# Characterization of binding kinetics and intracellular signaling of new psychoactive substances targeting cannabinoid receptor using transition-based reweighting method

**DOI:** 10.1101/2023.09.29.560261

**Authors:** Soumajit Dutta, Diwakar Shukla

## Abstract

New psychoactive substances (NPS) targeting cannabinoid receptor 1 pose a significant threat to society as recreational abusive drugs that have pronounced physiological side effects. These greater adverse effects compared to classical cannabinoids have been linked to the higher downstream *β*-arrestin signaling. Thus, understanding the mechanism of differential signaling will reveal important structure-activity relationship essential for identifying and potentially regulating NPS molecules. In this study, we simulate the slow (un)binding process of NPS MDMB-Fubinaca and classical cannabinoid HU-210 from CB_1_ using multi-ensemble simulation to decipher the effects of ligand binding dynamics on downstream signaling. The transition-based reweighing method is used for the estimation of transition rates and underlying thermodynamics of (un)binding processes of ligands with nanomolar affinities. Our analyses reveal major interaction differences with transmembrane TM7 between NPS and classical cannabinoids. A variational autoencoder-based approach, neural relational inference (NRI), is applied to assess the allosteric effects on intracellular regions attributable to variations in binding pocket interactions. NRI analysis indicate a heightened level of allosteric control of NPxxY motif for NPS-bound receptors, which contributes to the higher probability of formation of a crucial triad interaction (Y^7.53^-Y^5.58^-T^3.46^) necessary for stronger *β*-arrestin signaling. Hence, in this work, MD simulation, data-driven statistical methods, and deep learning point out the structural basis for the heightened physiological side effects associated with NPS, contributing to efforts aimed at mitigating their public health impact.

## Introduction

Cannabinoid receptor 1 (CB_1_), which is majorly expressed in the central nervous system (CNS) belongs to the class A G-protein coupled receptor (GPCR) family proteins. ^1–4^ GPCRs are expressed in the cellular membrane and help transduce chemical signals from the extracellular to the intracellular direction with the help of the downstream signaling proteins (G-proteins and *β*-arrestin). ^5–7^ In addition, GPCRs are the largest family of drug targets due to their substantial involvement in human pathophysiology and druggability.^8,9^ Significant research efforts have been invested in the discovery of drugs targeting CB_1_, which helps to maintain homeostasis in neuron signaling and physiological process.^10,11^

Initial drug discovery efforts, especially the design of synthetic agonists, were based on modifying the scaffolds of phytocannabinoids (e.g., Δ^9^-Tetrahydrocannabinol, cannabinol) and endocannabinoids (e.g., Anandamide, 2-arachidonoylglycerol) (Figure 1). ^12–14^ The synthetic molecules, which maintain the aromatic, pyran, and cyclohexenyl ring of the most common psychoactive phytocannabinoid Δ^9^-THC, are known as classical cannabinoids (Figure S1).^15–17^ However, the pharmacological potential of these molecules was diminished due to their psychological and physiological side effects (“tetrad” side effect).^18–20^ One such example of a synthetic cannabinoid is 1,1-Dimethylheptyl-11-hydroxy-tetrahydrocannabinol (commonly known as HU-210), which is a Schedule I controlled substance in the United States.^21,22^

**Figure 1.**
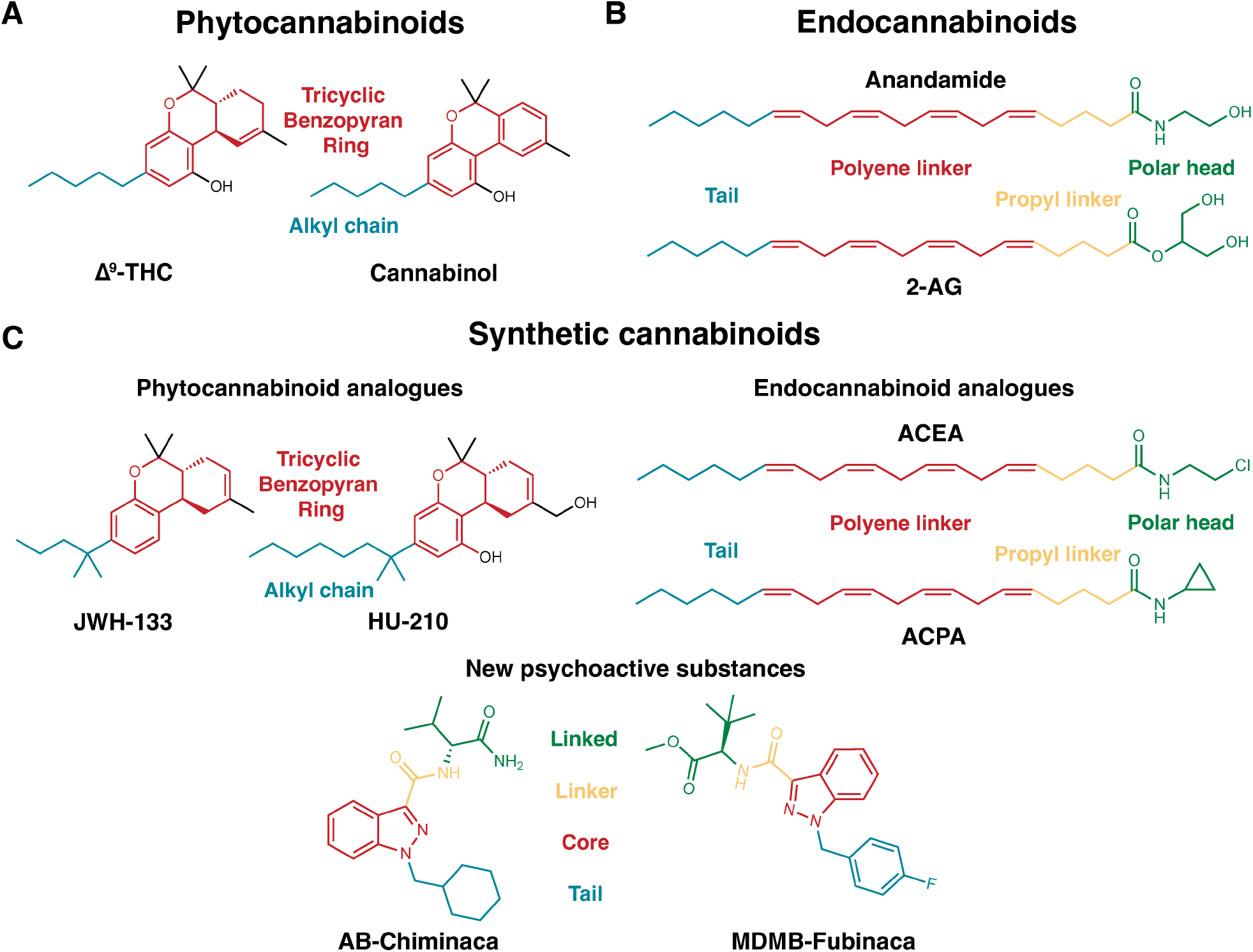
Classification of cannabinoid agonists: (A) Molecules derived from cannabis plants (phytocannabinoids) (B) endogenous agonists (Endocannabinoids) (C) synthetically designed molecules (Synthetic cannabinoids). Synthetic cannabinoids can be further classified based on scaffolds (phytocannabinoid analogues and endocannabinoid analogues or new psychoactive substances). Common pharmacophore groups of the ligands are shown in different colors. For phytocannabinoids and phytocannabinoid synthetic analogues, tricyclic benzopyran group and alkyl chains are colored in red and blue, respectively. Polar head group, propyl linker, polyene linker, and tail group of endocannabinoid and endocannabinoid analogues are colored with green, yellow, red, and orange, respectively. Linked, linker, core, and tail group of new psychoactive substances are colored with green, yellow, red, and orange, respectively.

Apart from the canonical structures of synthetic cannabinoids, molecules with diverse scaffolds were also synthesized through structure-activity studies. ^23–25^ However, these molecules also lacked any pharmacological importance due to psychological side effects.^26,27^ Due to the diverse structures and psychological effects, these molecules became unregulated substitutes for traditional illicit substances. ^28^ These synthetic cannabinoids belong to a class of molecules known as new psychoactive substances(NPS) as these molecules are not scheduled under the Single Convention on Narcotic Drugs (1961) or the Convention on Psychotropic Substances (1971).^28,29^ Synthetic cannabinoids make up the largest category of NPS molecules. ^30,31^ NPS creates a significant challenge for drug enforcement agencies, as they appeal to drug users seeking “legal highs” to avoid the legal consequences of using traditional drugs and to be undetectable in drug screenings.^27^

The molecular structures of NPS synthetic cannabinoids consist of four pharmacophore components: linked, linker, core, and tail groups. ^27,32^ The core usually consists of aromatic scaffolds (e.g., indole, indazole, Carbazole, Benzimidazole) (Figure S2). ^24^ The tail and linker groups are connected to the core. In the tail group, long alkyl chain-like scaffolds are ubiquitous in most NPSs; however, molecules with bulkier cyclic chains (e.g., AB-CHMINACA) are also present.^32^ Frequently encountered scaffolds in linker groups are methanone, ethanone, carboxamide, and carboxylate ester groups. ^33^ The linker acts as a bridge between the core and the linked group. In the initial NPS synthetic cannabinoids, the linked group included polyaromatic rings; however, non-cyclic linked groups have also been identified in NPS recently.^24,32^ Structural diversity in every component, while maintaining high binding affinity and potency for CB_1_ make these molecules easier for drug manufacturers and harder to ban by drug enforcement agencies. ^34,34–37^

The use of NPS synthetic cannabinoids has been found to cause more physiological side effects than traditional cannabinergic “tetrad” side effects. ^38^ These side effects include tachycardia, drowsiness, dizziness, hypertension, seizures, convulsions, nausea, high blood pressure, and chest pain.^38,39^ For instance, Gatch and Forster have shown that the high concentrations of AMB-FUBINACA, the molecule which caused “zoombie outbreak” in New York, induced tremors.^40,41^ A recent biochemical study has linked these discriminatory effects with the differential signaling of *β*-arrestin. ^39^ According to Finlay et al., NPS shows higher *β*-arrestin signaling compared to the classical cannabinoids, which has also been confirmed by other *β*-arrestin signaling studies. ^39,42^ However, mechanistic understanding of these differential downstream signaling effects between NPS and classical cannabinoids is still missing.

Mutagenesis studies have shown that the conserved NPxxY motif of CB_1_ have a larger role in downstream *β*-arrestin signaling than G-protein signaling. ^43,44^ Recently published MDMB-FUBINACA bound CB_1_-*β*-arrestin-1 complex structure also points out the importance of the unique triad interaction (Y397^7.53^-Y294^5.58^-T210^3.46^) involving NPxxY motif in *β*-arrestin-1 signaling. ^44^ However, structural comparison of the classical cannabinoid (AM841) and NPS (MDMB-FUBINACA) bound active CB_1_-G_*i*_ complex shows a conformationally similar NPxxY motif (Figure 2).^45,46^ In light of these experimental observations, it can be inferred that higher *β*-arrestin signaling stems from higher dynamic propensity of triad interaction formation for NPS-bound CB_1_. We hypothesized that distinct orthosteric pocket interactions for NPS and Classical Cannabinoids causes differential allosteric modulation of intracellular dynamics that facilitate triad interaction.

**Figure 2.**
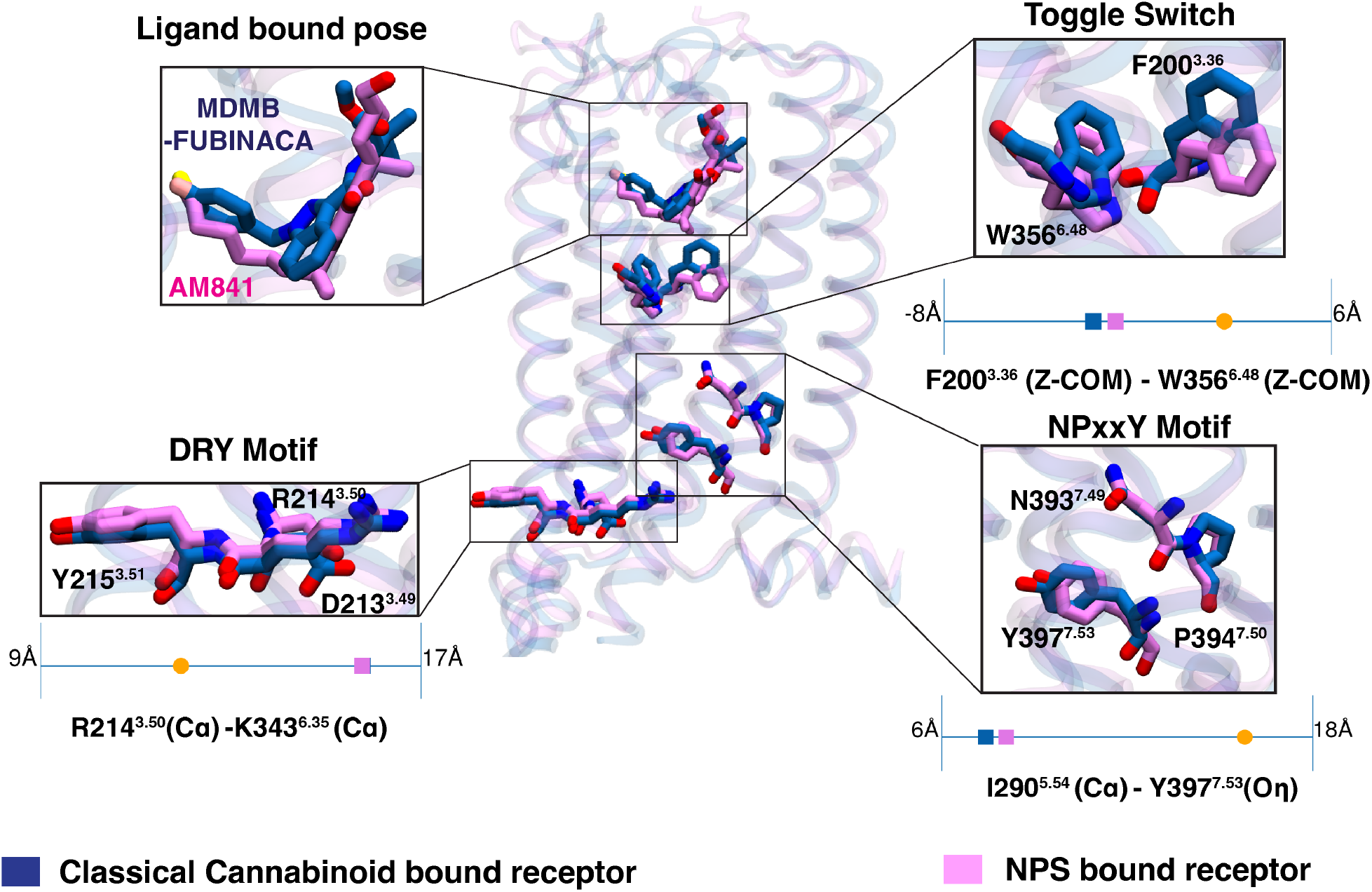
NPS bound CB_1_ (PDB ID: 6N4B, ^45^ color: Blue) structure is superposed with the classical cannabinoid bound CB_1_ (PDB ID: 6KPG, ^46^ color: Purple). Both structures are in G_*i*_ bound active state. Proteins are shown in transparent cartoon representation. Structural comparison of conversed activation metrices (Toggle switch, DRY motif, and NPxxY motif) and ligand poses are shown as separate boxes. Quantitative values of the activation metrics for both active structures are compared as scatter points on 1-D line with the CB_1_ inactive structure (PDB ID: 5TGZ, ^1^ color: orange). These quantitative measurements were discussed in Dutta and Shukla ^4^

To study these distinct dynamic effects, we compared the (un)binding of the classical cannabinoid (HU-210) and NPS (MDMB-FUBINACA) from the receptor binding site. These molecules have nanomolar affinities. Obtaining the initial pathway of ligand unbinding from unbiased sampling will be computationally expensive. Therefore, a well-tempered metadynamics approach was used to sample the unbinding event, where a time-dependent biased potential is deposited for the faster sampling of the metastable minima along the pathway.^47^ However, a detailed characterization of the unbinding processes is only possible through the thermodynamics and kinetics estimation of intermediate states. Thus, a transition operator-based approach is needed, which helps to estimate the transition timescale between the states and the stationary density of each state. Estimation from these approaches usually depends on the equilibrium between the local states, which can only be maintained by reversible sampling. For high-affinity ligands like MDMB-FUBINACA and HU-210, reversible sampling is expensive as ligands move from high energy unbound states to lower energy bound states irreversibly. Hence, we implemented a transition operator approach named the transition-based reweighting analysis (TRAM) method, which can tackle this lack of local equilibrium between states by combining unbiased and biased approaches. ^48^ TRAM has been used different simulation studies for estimating thermodynamics and kinetics of processes that have high free energy barriers. For example, TRAM have been utilized for characterization of small molecule and peptide (un)binding processes,^48–52^ protein dimerization,^53^ ion transportation.^54^ To implement TRAM for our study, extensive sampling of the (un)binding process of both ligands was performed using a combination of umbrella sampling and unbiased simulations from the pathway obtained using metadynamics (see Methods section).^55^ We showed that TRAM can produce consistent kinetic estimation with less unbiased simulation data compared to traditional methods like the Markov state model. ^56^

Based on estimates of thermodynamics and kinetics, it was observed that both NPS and classical cannabinoids have similar unbinding pathways. However, their unbinding mechanisms differ due to the aromatic tail of the MDMB-FUBINACA compared to the alkyl side chain of HU-210. Furthermore, dynamic interaction calculations reveal a major difference with TM7 between NPS and classical cannabinoid. Specifically, the hydroxyl group in the benzopyran moiety of HU-210 forms much stronger polar interactions with S383^7.39^ compared to the carbonyl oxygen of the linker group in MDMB-FUBINACA. MD simulations of other classical cannabinoids and NPS molecules bound CB_1_ also support these significant interaction differences. The ligand binding effect in intracellular signaling was estimated by measuring the probability of triad formation in the intracellular region. NPS bound CB_1_ shows higher probability of forming triad interaction compared to the classical cannbinoids, which supports the experimental observations of high *β*-arrestin signaling of NPS bound receptors. To validate that the triad formation is indeed caused by the binding pocket interaction differences between the two ligands, allosteric strength binding pocket residues and NPxxY motif was estimated with the deep learning technique, Neural relational inference (NRI).^57^ NRI network revealed that binding pocket residues of NPS bound ensemble have higher allosteric weights for the NPxxY motif compared to classical cannabinoids. These analyses validate our hypothesis that the differential dynamic allosteric control of the NPxxY motif might lead to the *β*-arrestin signaling for different ligands. This study provides a foundation for additional computational and experimental research to enhance our understanding of the connection between ligand scaffolds and downstream signaling. This knowledge will assist drug enforcement agencies in proactively banning these molecules and inform policies that can protect individuals from the effects of abuse.

## Results and Discussion

### Metadynamics simulations capture the unbinding paths of NPS and classical cannabinoids

The representative classical cannabinoid and NPS selected for this study are HU-210 and MDMB-FUBINACA.^22^ Compared to Δ^9^-THC, HU-210 has an extra hydroxyl group in the C-11 position and a 1’,1’-Dimethylheptyl group instead of a pentyl side chain (Figure 1C and S1). MDMB-FUBINACA is a derivative of AB-FUBINACA, which was originally developed by Pfizer (Figure 1C).^45^ These ligands binds to CB_1_ receptor with nanomolar affinities (MDMB-FUBINACA K_*i*_ : 1.14 nM;^58^ HU-210 K_*i*_ : 0.61 nM^14,59^).

Metadynamics simulation is a biased sampling method and has been widely used in protein-ligand binding and unbinding studies, as preexisting knowledge of the pathway is not necessary for performing these simulations.^60–63^ In metadynamics, a time-dependent biased potential is deposited into the sampling process for the ligand to get out of stable minima at a faster pace.^64^ In this work, two replicates of well-tempered metadynamics were performed to capture the unbinding pathway of HU-210 and MDMB-Fubinaca. The commonly used collective variables were selected for metadynamics simulations: (1) z component distance between the center of mass of ligand and residue in the ligand binding pocket (W356^6.48^), and (2) Contact number with the ligand heavy atom and *α* carbon of all binding pocket residues (Equation 4).

The z-component distance was plotted against the RMSD of the ligands from the bound pose, which indicates that ligands follow a similar pathway for each replica (Figures S3A and S3C). It is observed that the dissociation happens via the opening formed by TM2, TM3, ECL2, and N-terminus for both ligands (Figures 3A, 3B, 3C, and 3D). We also performed unbinding simulations using well-tempered metadynamics parameters (bias height, bias deposition rate and biasfactor) to confirm the existence of alternative pathways (Figure S4). However, the simulations show that ligands follow the similar pathway for all metadynamics runs. These observations indicate that the pathway may be the minimum free energy pathway for the ligand unbinding in CB_1_. Previous metadynamics binding simulation of another cannabinoid ligand also points to a similar pathway. ^63^ Reweighted probability density obtained from metadynamics calculation shows one highly dense region in the pocket, depicting the stability of the bound pose of the ligands (Figures S3C and S3D). However, time dependent external force applied during the metadynamics makes the sampling in the orthogonal direction of the CVs less extensive. Thus, the biased simulation might not sample some protein-ligand interactions that helps to characterize intermediate states. To properly characterize intermediate transition states during the unbinding process, discrete kinetic models based on extensive unbiased simulations have been used. These unbiased simulations are often initiated from the pathways derived from the initial, limited sampling obtained through biased simulations^49,65,66^ (discussed below).

**Figure 3.**
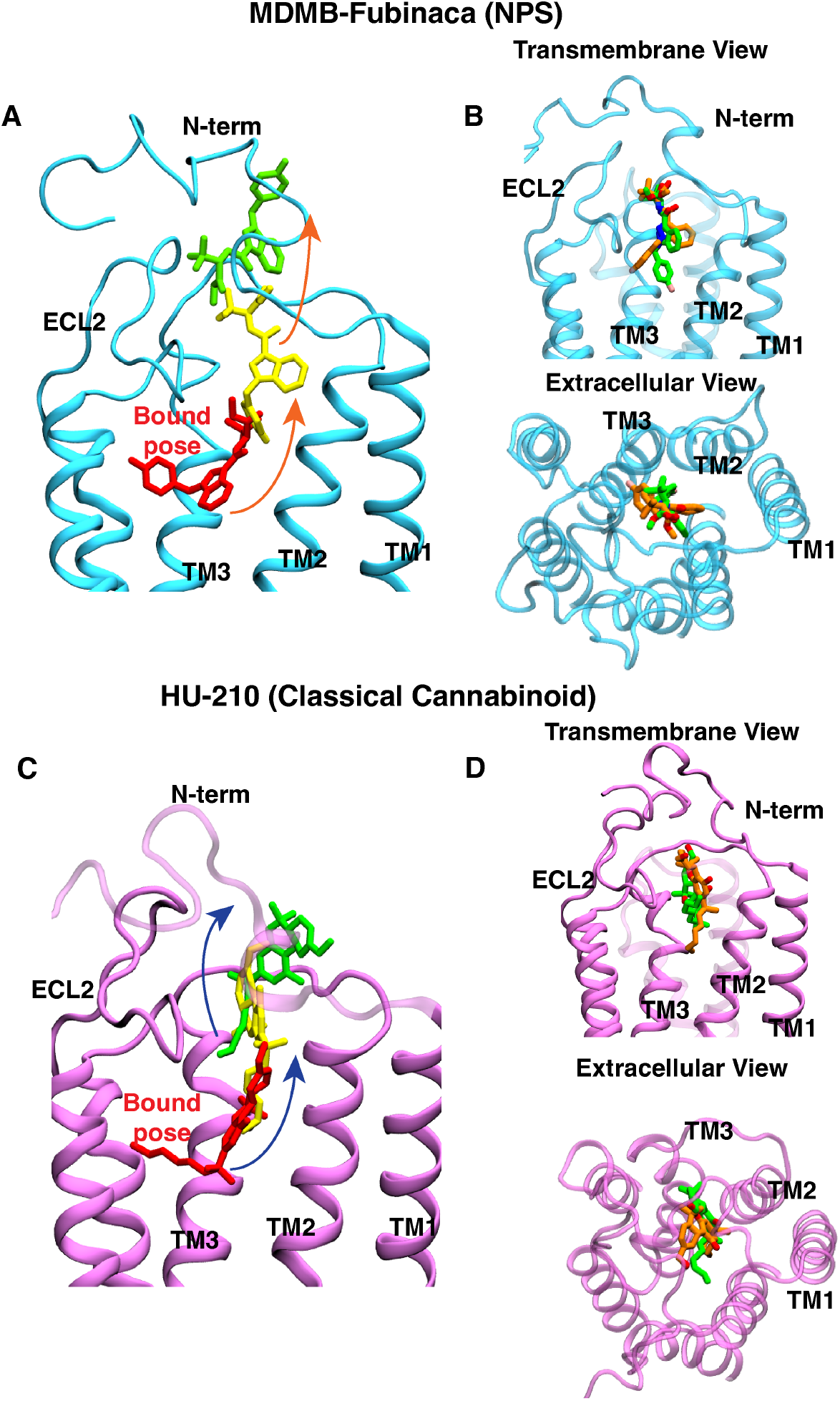
The ligands MDMB-FUBINACA (A) and HU-210 (C) are depicted in three distinct stages along their unbinding pathways, as determined by well-tempered metadynamics simulations. The ligands are illustrated using stick representations, with each stage represented by a different color to indicate the progression from the bound (color: red) to the unbound state (color: green). Representative ligand positions from an intermediate state are shown in yellow. Additionally, the superposition of representative frames of an intermediate stage of the unbinding process is shown, where MDMB-FUBINACA (B) and HU-210 (D) are dissociating from the receptor. The frames are obtained from two different well-tempered metadynamics simulation replicas and are shown with different colors (green and orange). Both transmembrane (left panel) and extracellular (right panel) views are displayed. Proteins are represented as cartoons.

### Comparison of thermodynamics and kinetics estimates from Markov state Model and Transition-based reweighting analysis method

MSM and TRAM are both postprocessing techniques for estimating the kinetics and thermodynamics of underlying physical processes observed in MD simulation. MSM is applied to reversible equilibrium simulations, whereas TRAM estimations can be obtained from multi-ensemble simulations (combination of biased and unbiased simulations). The MSM depends on the local equilibrium between the Markovian states, which is also known as detailed balance. However, reversible local sampling becomes challenging with short parallel trajectories when the free energy difference between two local Markovian states is high. In those cases, reversibility is still assumed by forcing the detailed balance when estimating the transition probability matrix.^56^ This leads to the incorrect estimation of the unbinding kinetics due to limited sampling from the stable bound state to the high energy unbound states. ^48^ Refining the state discretization (i.e., increasing the number of states) may resolve the issue. However, refined state discretization sometimes decreases the statistically significant transition count between all states, decreasing the model certainty. TRAM was shown to solve this problem by combining biased and unbiased simulations (see Methods section). Biased simulations (e.g., replica exchange, umbrella sampling) help to enhance the local sampling, either by increasing the temperature for faster sampling or by fixing collective variables with biased potential to have better sampling in orthogonal directions. It has been shown that compared to MSM, kinetics predicted using TRAM from the combination of biased and unbiased simulations are more aligned with the experiment results. ^48^

As unbinding of ligands with high binding affinity (nanomolar) are being studied here, asymmetric transitions might be observed along the pathway. Therefore, we compared the use of MSM and TRAM in estimating the kinetics and thermodynamics of the (un)binding process. For TRAM, unbiased simulation and umbrella sampling were run from the clusters in conformational ensemble obtained from metadynamics (refer to the Methods section for more details). For MSM estimation, only unbiased simulations starting from the metadynamics pathway were considered.

For thermodynamics comparison, standard free energy was estimated for the ligands considering volume correction. ^67^ TRAM and MSM predictions of standard binding free energy are within 0.6 kcal/mol of each other for each ligand (Figure 4A). Although the absolute binding free energy differs from the experimentally predicted value by approximately 3 kcal/mol, the relative estimated free energy (ΔΔG) values are also within 0.6 kcal/mol of the experimentally determined values. Therefore, it indicates that with sufficient sampling, both MSM and TRAM converge to the same predictions of relative free energy.

**Figure 4.**
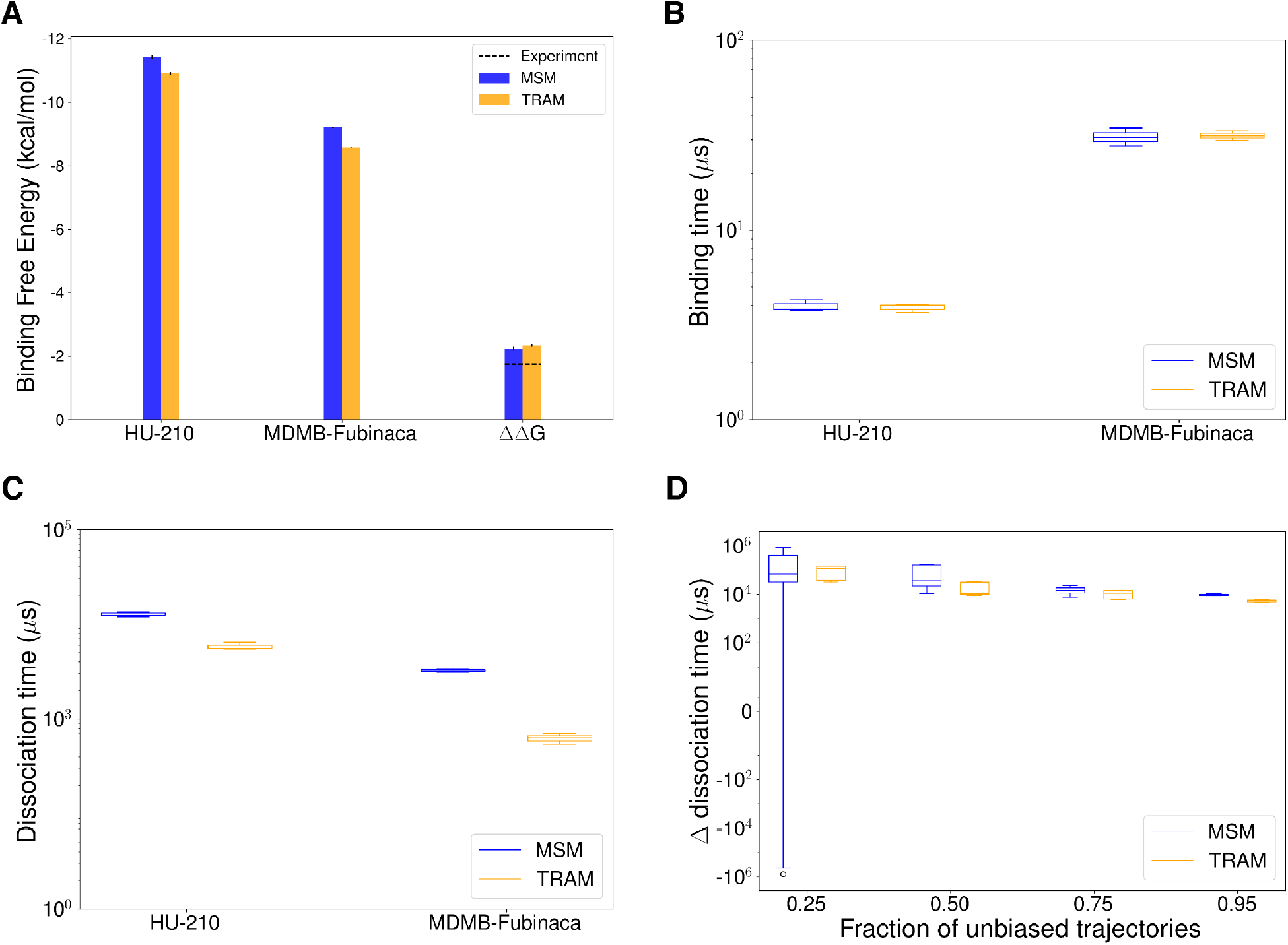
(A) The bar plot represents standard binding free energy for HU-210, MDMB-FUBINACA, and difference of standard binding free energy between the ligands. MSM and TRAM estimations are shown as blue and orange bars, respectively. Experimentally predicted values are shown as dotted line. (B, C) Binding (B) and dissociation (C) time for HU-210 and MDMB-FUBINACA are shown as box plots. (D) Difference in dissociation time of the two ligands is plotted as box plot against fraction of unbiased trajectories used for the estimation. This timescales were obtained from the mean free passage time calculation using TPT with transition probabilities estimated from MSM (color: blue) and TRAM (color: orange). Errors were calculated using bootstrapping method with 3 bootstrapped samples.

We also compared the kinetics obtained from the MSM and TRAM. Kinetic measurements were performed with transition path theory (TPT), which uses transition probability matrix from MSM or TRAM to estimate mean free passage time between different states (see Methods section). Estimated binding times using TRAM and MSM match perfectly for both ligands. The estimated dissociation times are within one order of magnitude with each other. These observations agree with the previously reported computational research, where experimentally comparable estimation of *k*_*off*_ rates were shown to be more challenging compared to *k*_*on*_.^68^

Further analyses were performed to compare these methods in the low unbiased data regime. The difference between the dissociation time of ligands was measured with different amounts of unbiased data. It is observed that even with only 25% of original, unbiased data, TRAM can predict the kinetics within an order magnitude of the kinetics estimated with full dataset (Figure 4D). On the other hand, error in MSM predicted kinetics more rapidly compared to TRAM with lesser amount of unbiased data. A similar trend can be observed for ΔΔG prediction (Figure S5). Therefore, TRAM provides better predictions of thermodynamics and kinetics when less amount of unbiased data.

### Unbinding mechanism for new psychoactive substance

Although the binding position of the ligand and the overall binding pathway are similar for both the ligands, extensive biased and unbiased simulation analyzed by TRAM shows a significant difference in the unbinding mechanism of the ligands. To capture the unbinding pathway for MDMB-FUBINACA, we projected the TRAM weighted free energy landscape of the distance between the linked part of the ligand (Leucinate group) and TM5 with respect to the distance between the ligand tail group and TM7 (Figure 5A). The free energy landscape was divided into the non-overlapping intermediate macrostates to obtain better description of the unbinding process. In each macrostate, the contact frequency of ligand with binding pocket residues were calculated along with corresponding contact energies. A metastable minima is observed for macrostate representing the bound pose of the ligand depicting the stability of the ligand (Figure 5A). In the bound pose, the major interactions form between the aromatic (F170^2.57^, W279^5.43^, F268^ECL2^) and hydrophobic (L193^3.29^) residues of the binding pocket (Figures 6A, 6B, and S6B).

**Figure 5.**
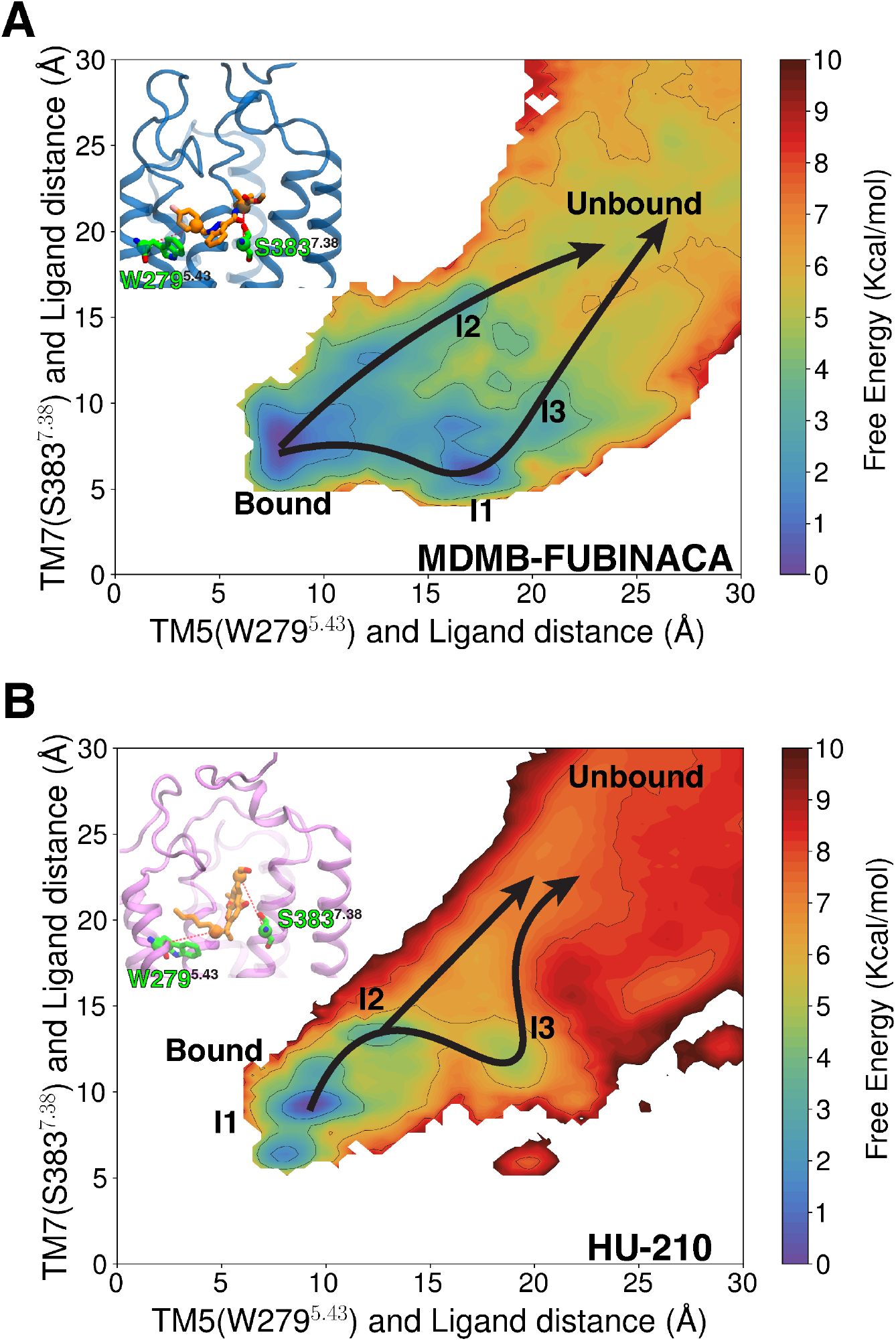
TRAM weighted Two dimensional projection of unbinding free energy landscape for MDMB-FUBINACA (A) and HU-210 (B). For MDMB-FUBINACA, distance between TM5 (W279^5.43^-C*α*) and tail part of the ligand is plotted against the distance between TM7 (S383^7.39^-C*α*) and ligand linked part. For HU-210, distance between the TM5 (W279^5.43^-C*α*) and tail is plotted against the TM7 (S383^7.39^-C*α*) and cyclohexenyl ring of the ligand. Measured distances are shown as red dotted lines in the inset figures. Macrostate positions are shown on the landscapes. Different mechanisms of (un)binding are shown with arrow on top of the free energy landspace.

**Figure 6.**
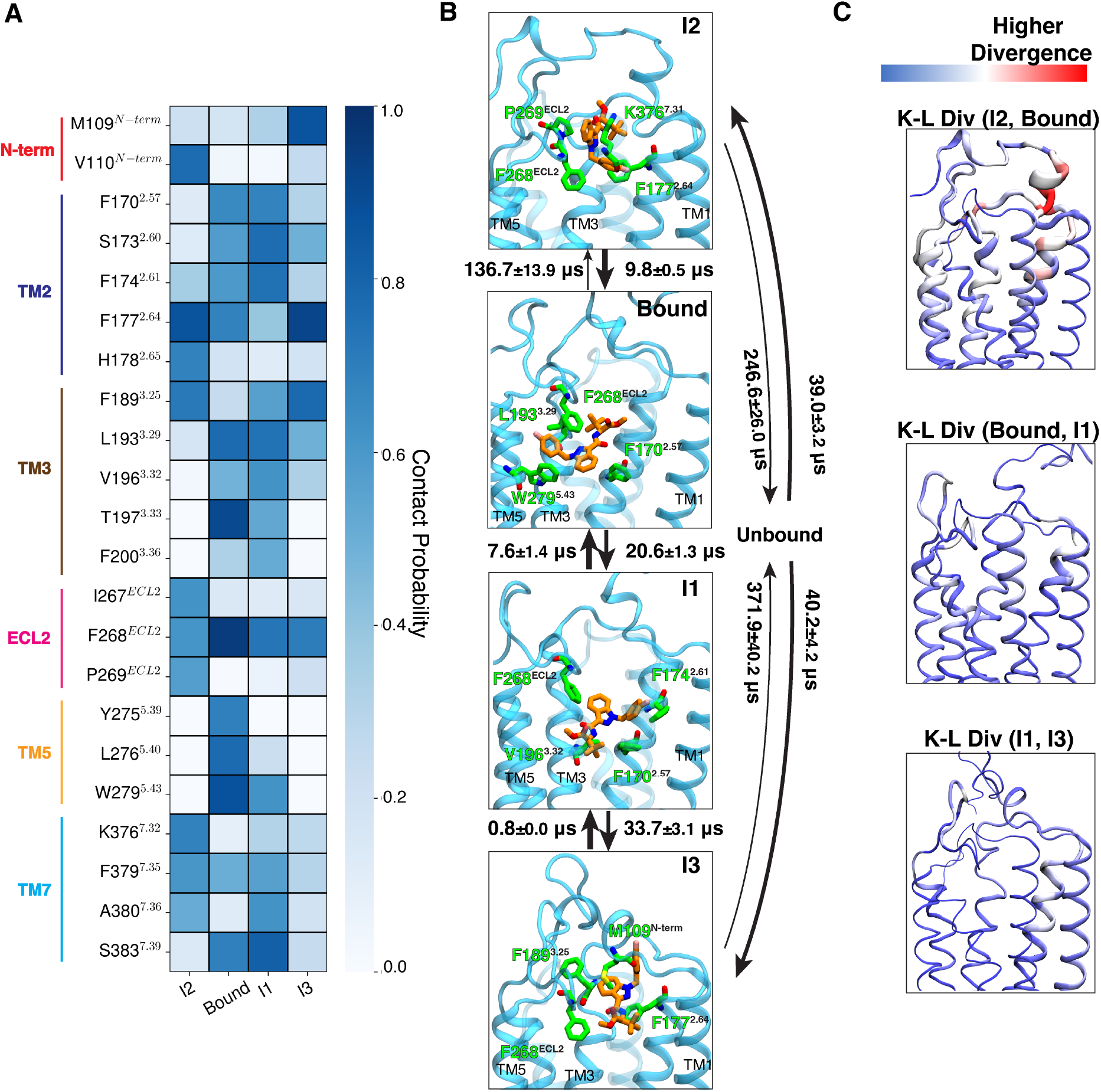
(A) The contact probabilities with binding pocket residues of MDMB-FUBINACA are shown as a heatmap for different macrostates, where ligand maintains contact with the receptor. Residues in different structural elements (loops and helices) are distinguished by distinct color bars. (B) Representative structures are shown where ligand (color: orange) and four residues (color: green) with highest interaction energies are shown as sticks. Proteins are shown as purple cartoon. Timescales between interstate transitions are shown as arrows. Arrow thickness is inversely proportional to the order of magnitude of the timescale. (C) Per residue K-L divergences between different states are shown with color (blue to red) and thickness (lower to higher) gradient. K-L divergences calculated on the inverse distance feature distributions were converted by residue basis by summing all the pair contributions corresponding to the residue. Thickness gradients are shown as rolling average to highlight a region of high K-L divergence. Errors in MFPT calculations were estimated based on 3 bootstrapped TRAM calculation with randomly selected 95% of unbiased trajectories.

The free energy landscape shows two probable mechanisms for MDMB-FUBINACA unbinding from the bound pose. The two pathways are differentiated by whether the linked or tail part of MDMB-FUBINACA dissociates first. One of the pathways, aromatic tail part of MDMB-FUBINACA moves away from TM5 and form interaction with aromatic residues in TM2 (F170^2.57^ and F174^2.61^) (Figures 5A, 6A, 6B, and S6C). This leads to the formation of intermediate metastable states, which we characterize as macrostate Intermediate state 1 (I1). This metastable minimum observed from I1 macrostate might be unique to the FUBINACA family of NPS synthetic cannabinoids as this family has the aromatic ring in tail group, unlike the long alkyl chain in other common synthetic cannabinoids. Along with the aromatic residues of TM2, major interaction with F268^ECL2^ is maintained in macrostate I1 (Figures 6A and 6B). KL divergence analysis of the inverse distance distributions between two macrostates was conducted to highlight significant conformational changes. of bound and I1 macrostates show that only minor changes in the binding pocket residues, especially in TM2 are needed to accommodate MDMB-FUBINACA in this conformational state (Figure 6C). The interconversion timescale (MFPT) between the macrostates were obtained from the transition path theory. MFPT calculations show that both the timescales are similar with slightly higher timescales for the bound pose compared to I1 transition (20.6 ± 2.3*µs*) (Figure 6B). In this pathway, ligand moves from I1 metastable state to space between N-terminus, TM2, TM3, and ECL2 before dissociating from the receptor (Figure 6B). This region between the unbinding ensemble has been characterized as macrostate I3 (Figure 5A). Contact analysis show significant drop in ligand residue contacts with only aromatic residues in TM2, TM3 and ECL2 forming dominant interactions (Figures 6A and S6D). We further performed Kullback-Leibler divergence (K-L divergence) analysis between inverse distance of residue pairs of two macrostates to highlight the protein region that undergoes high conformational change with ligand movement (detailed discussion in Methods section). K-L divergence shows that ligand positioning in this particular regions causes relatively higher divergence on TM2 compared to I1 (Figure 6C). Kinetically, the transition from I1 to I3 (33.7 ± 3.1*µs*) is much slower compared to reverse transition (0.8 ± 0.0*µs*), validating the higher stability of the I1 compared to I3 macrostate (Figure 6B). According to the TPT analysis, breaking the aromatic interactions for complete dissociation of MDMB-FUBINACA requires ∼ 371.9 ± 40.2*µs*, making it the slowest step in this pathway (Figure 6B).

In the other possible unbinding pathway, orientation of MDMB-FUBINACA in the pocket doesnot change compared to the bound pose. The linked part of the ligand moves to space between N-terminus, TM2, TM3, and ECL2 (Figure 6B). We label this macrostate as I2. In this state, we observe stable polar interaction with K376^7.32^ and hydrophobic interactions with aromatic and other hydrophobic residues (F177^2.64^, F268^ECL2^, P269^ECL2^) (Figures 6A and S6A). However, free energy of this macrostate is higher than the bound pose, depicting higher entropic cost associated with this state. This can be shown by the higher intrastate RMSD of I3 compared to the bound pose (Figure S7). Transition timescale from the bound pose to the I2 (136.7 ± 13.9*µs*) is one order of magnitude higher compared to the reverse transition (9.8 ± 13.9*µs*) (Figure 6B). K-L divergence analysis also shows higher divergence in the extracellular region of TM2 and N-terminus compared to bound pose (Figure 6C). Dissociation of MDMB-FUBINACA from I2 to the bulk is faster compared to dissociation from I3 (246.6 ± 26.0*µs*) (Figure 6B). However, the overall kinetic barrier for dissociation from the binding pose for the both unbinding mechanisms are relatively similar.

### Unbinding mechanism for classical cannabinoid

For capturing the classical cannabinoid (un)binding mechanism, distances from the two terminal scaffolds (cyclohexenyl and alkyl chain) to TM5 and TM7 were measured similar to the NPS (Figure 5B). The free energy landscape of the unbinding of the HU-210 shows the differences in the mechanism from MDMB-FUBINACA. Similar to MDMB-FUBINACA, the HU-210 unbinding landscapes were also divided into non-overlapping macrostates. Macrostate representing HU-210 bound pose shows a metastable energy minimum. Comparing the bound macrostate interactions of MDMB-FUBINACA, classical cannabinoid HU-210 shows higher interactions with TM7 residues (S383^7.39^, F379^7.35^) (Figures 6A, 7A, S6B and S8B). Previous experimentally determined structures of classical cannabinoid bound CB_1_ have pointed out these conserved polar interactions of the hydroxyl group at C-1 position with S383^7.39^.^46,69^ Although MDMB-FUBINACA also maintain this polar interaction with carboxylic oxygen, the interaction energy for the HU-210 is much higher, depicting the importance of this conserved residue in stabilizing classical cannabinoids (Figure 6B). Mutagenesis studies also support this difference in interaction with S383^7.39^ between the classical cannabinoids with hydroxyl group (HU-210) and CB_1_ ligands, which have carboxylic oxygen in the equivalent position (WIN-55,212-2).^70,71^ Alanine mutation of S383^7.39^ have shown to decrease the ligand affinity and downstream efficacy of classical cannabinoids by orders of magnitude, while having minimal effect on WIN-55,212-2, which have carboxylic oxygen in the linked part as MDMB-FUBINACA.^70,71^ Other major interactions (F170^2.57^ and F268^ECL2^) in the bound pose are common between the two ligands (Figures 6B and 7B).

A relatively weaker metastable state is observed when the ligand moves relatively deeper (closer to TM5) inside the binding pocket. The flexible alkyl chain of HU-210 allows the ligand to have this deeper position (Figures 7A and 7B). Protein-ligand interaction analysis in the macrostate representing this region (I1) show that hydroxyl group at C-11 forms a major polar interaction with H178^2.65^ (Figure S8A). The bound and I1 macrostates are kinetically close, as indicated by the rapid interconversion between these states (Figure 7B). K-L divergence between the two states show the highest divergence in extracellular TM2 and TM7, where major interaction switch have happened (Figure 7C).

**Figure 7.**
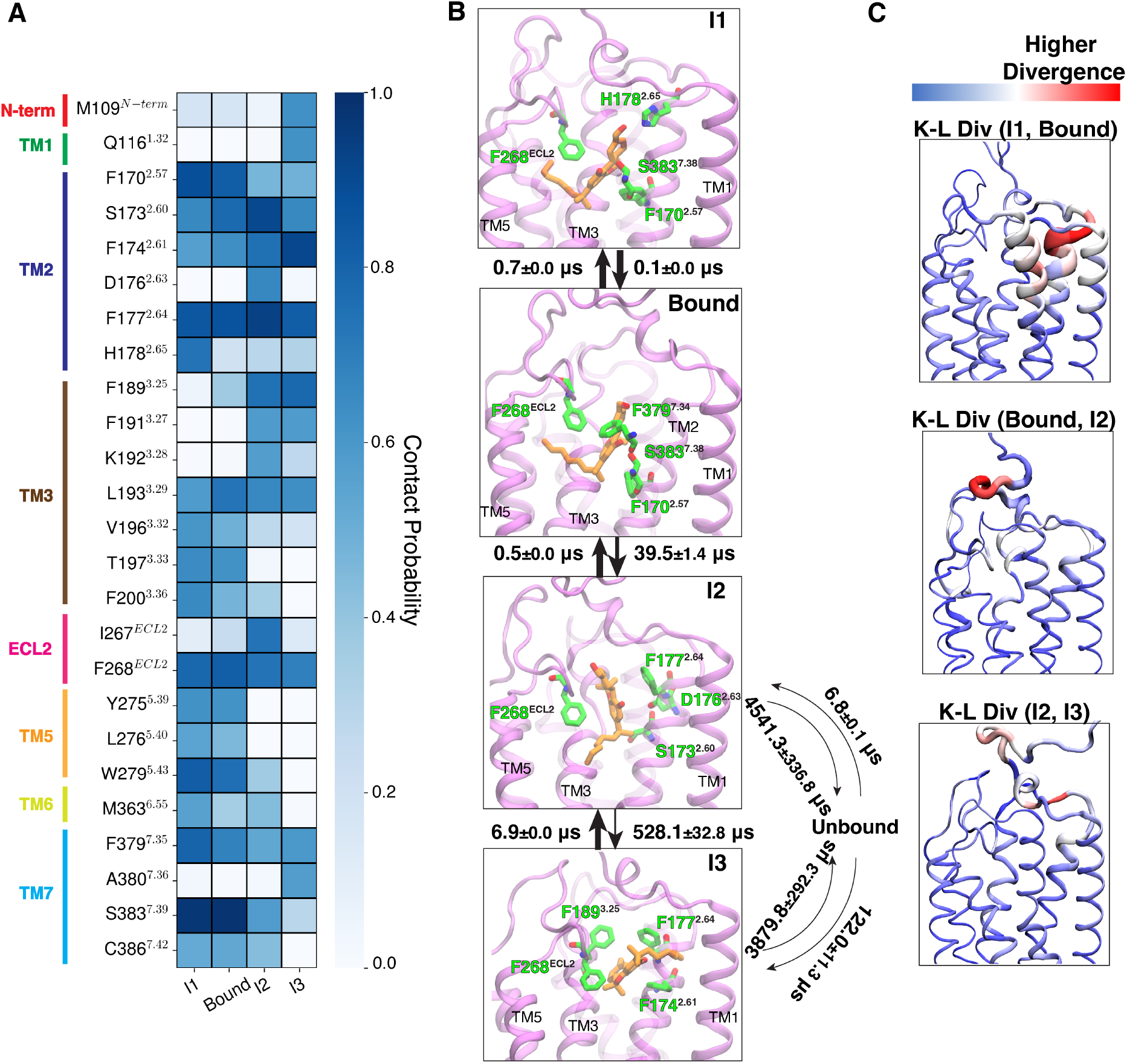
(A) The contact probabilities with binding pocket residues of HU-210 are shown as a heatmap for different macrostates, where ligand maintains contact with the receptor. Residues in different structural elements (loops and helices) are distinguished by distinct color bars. (B) Representative structures are shown where ligand (color: orange) and four residues (color: green) with highest interaction energies are shown as sticks. Proteins are shown as purple cartoon. Timescales between interstate transitions are shown as arrows. Arrow thickness is inversely proportional to the order of magnitude of the timescale. Errors in MFPT calculations were estimated based on 3 bootstrapped TRAM calculation with randomly selected 95% of unbiased trajectories. (C) Per residue K-L divergences between different states are shown with color (blue to red) and thickness (lower to higher) gradient. K-L divergences calculated on the inverse distance feature distributions were converted by residue basis by summing all the pair contributions corresponding to the residue. Thickness gradient are shown as rolling average to highlight a region of high K-L divergence.

Contrasting to MDMB-FUBINACA, only one pathway was discovered with classical cannabinoid cyclic scaffold departing from the receptor first. Major interactions that break when the ligand moves out of the binding pose to macrostate I2 is the polar interaction with S383^7.39^ and hydrophobic interaction with aromatic F170^2.57^ (Figures 7A, 7B, and S8C). Breaking of these bonds leads to larger kinetic barrier of approximately 39.5 ± 1.4*µs* (Figure 7B). In this macrostate, the HU-210 forms major interactions with aromatic residues F268^ECL2^ and F177^2.64^ and polar interactions with S173^2.60^ and D176^2.63^ (Figure 7B). From this pose, HU-210 either dissociates from the receptor or obtain another relatively weak stabilized state (I3) in the receptor. In I3, the alkyl chain of the ligand is flipped in the pocket and stabilized by aromatic residues in TM2, TM3 and ECL2 (Figure 7B). This transition from I2 to I3 (528.1 ± 32.8*µs*) kinetically much slower compared to the reverse transition (6.9 ± 0.0*µs*) (Figure 7B). From both I2 and I3 macrostates, the ligand can dissociate from the pocket and mean free passage time for these transitions appear to be in the milisecond timescale, which is one order magnitude higher compared to the MDMB-FUBINACA unbinding (Figure 6B). This phenomena supports the relatively high affinity of the classic cannabinoid HU-210 compared to the NPS MDMB-FUBINACA.

### Allosterically controlled distinct downstream signaling between new psychoactive substances and classical cannabinoids

As discussed in the previous section, major interaction differences between NPS MDMB-FUBINACA and classical cannbinoid HU-210 are observed in TM7. To support the universality of this observation, we performed unbiased MD simulation (1 *µ*s each) of other NPS (AMB-FUBINACA, 5F-AMP, CUMYL-FUBINACA) and classical cannabinoids (AMG-41, JWH-133, O-1317) bound CB_1_ (Figure S9A). Average distance of carbonyl oxygen of NPS molecules’ linker group from S383^7.39^ is compared to equivalent distance of hydroxyl group of classical cannbinoids’ benzopyran ring. Larger mean distance in case of all NPS-bound CB_1_ supports the universality of the weaker interaction between TM7 and NPS molecules (Figure S9B). This variation in binding pocket pocket interactions might lead to differential allosteric control of the intracellular dynamics that facilitate triad interaction (Y397^7.53^-Y294^5.58^-T210^3.46^) important for *β*-arrestin binding.

We adopted a data-driven deep learning network known as Neural relational inference (NRI) to validate our hypothesis of allosteric control. NRI network has an architecture of variational autoencoder. The encoder part of the network predicts the interactions between the residues from the trajectory dynamics, and the decoder predicts the trajectories from the interaction. With this network, we try to produce alpha carbon coordinates at *t* + *τ* from the coordinates at time t. In the process of regenerating the future coordinates, the latent space of the network learns the dynamic interactions between different residues in the protein.

These interactions are calculated from the estimated posterior probability *q*(*z*_*ij*_|*x*). In this work, we trained the network with the NPS (MDMB-FUBINACA), and classical cannabinoid (HU-210) bound unbiased trajectories (Method Section). Here, we compared the allosteric interaction weights between the binding pocket and the NPxxY motif which involves in triad interaction formation. Results show that each binding pocket residue in MDMB-FUBINACA bound ensemble shows higher allosteric weights with the NPxxY motif, indicating larger dynamic interactions between the NPxxY motif and binding pocket residues(Figure S10). To further validate our observations, we estimated allosteric weights between the binding pocket and the NPxxY motif by calculating mutual information between residue movements. Mutual information analysis reaffirms that allosteric weights between these residues are indeed higher for the MDMB-FUBINACA bound ensemble (Figure S11).

The probability of triad formation was estimated to observe the effect of the difference in allosteric control. TRAM weighted probability calculation showed that MDMB-FUBINACA bound CB_1_ has the higher probability of triad formation (Figure 8A). Comparison of the pairwise interaction of the triad residues shows that interaction between Y397^7.53^-T210^3.46^ is relatively more stable in case of MDMB-FUBINACA bound CB_1_, while other two interactions have similar behavior for both systems (Figures S12A, S12B, and S12C). Therefore, higher interaction between Y397^7.53^ and T210^3.46^ in MDMB-FUBINACA bound receptor causes the triad interaction to be more probable.

**Figure 8.**
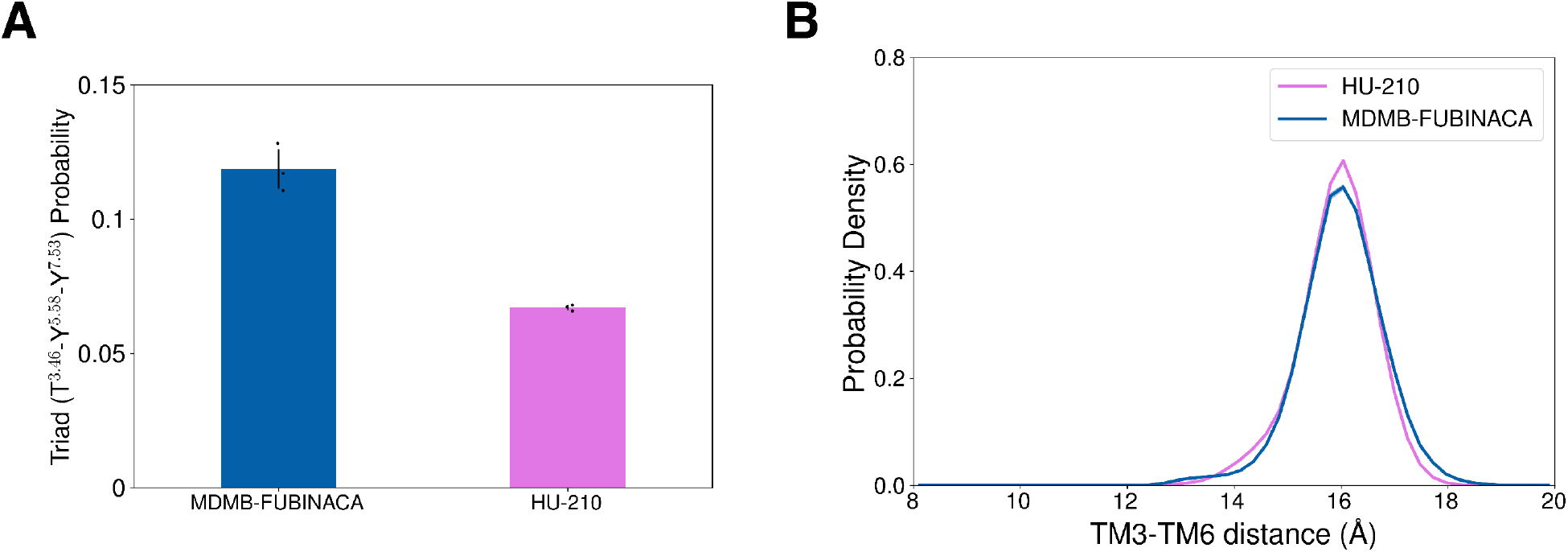
(A) TRAM weighted probabilities of triad interaction (Y397^7.53^-Y294^5.58^-T210^3.46^) formation are plotted for HU-210 (color: purple) and MDMB-FUBINACA (color: blue) unbinding ensemble. If side-chain oxygen atoms of all three residues are within 5 Å of each other, triad interaction is considered to be formed. (B) TRAM weighted probability densities of TM3 (R214^3.50^) and TM6 (K343^6.35^) distance distribution are plotted for HU-210 (color: purple) and MDMB-FUBINACA (color: blue) unbinding ensemble. Error in the probability densities is estimated using bootstrapping approach, where TRAM was built for 3 bootstrapped samples with 95% of total data.

Furthermore, we also compared TM6 movement for both ligand bound ensemble which is another activation metric involved in both G-protein and *β*-arrestin binding. Comparison of TM6 distance from the DRY motif of TM3 shows similar distribution for HU-210 and MDMB-FUBINACA (Figure 8B). These observations support that NPS binding causes higher *β*-arrestin signaling by allosterically controlling triad interaction formation.

## Conclusions

Synthetic cannabinoids were designed as a potential therapeutics to target cannabinoid receptors. However, major side effects of these ligands diminish their therapeutic potential. Although both classical cannabinoids and NPS synthetic cannabinoids have been abused as recreational drugs, later poses larger threats for the society due the chemical diversity of the NPS structures makes them harder to control from being abused. Furthermore, physiological studies have shown NPS targeting cannabinoid receptors lead to the dangerous physiological effects compared to “tetrad” side effects associated with classical cannabinoids. Previous studies have relate these side effects with the higher downstream *β*-arrestin signaling of NPS. Mutagenesis studies have shown NPxxY motif have larger role to play in *β*-arrestin signaling. In this work, we proposed that NPS and classical cannabinoid have distinct allosteric control on NPxxY motif when bound to orthosteric pocket of CB_1_. In this hypothesis driven study, we compared ligand-protein interactions of NPS MDMB-FUBINACA and classical cannabinoid HU-210 for CB_1_ by studying their unbinding mechanism and downstream signaling.

As both ligands are stable binders with nanomolar affinity, well-tempered metadynamics simulations were performed to obtain the initial unbinding pathway. These simulations were able to find similar pathway via the opening formed by TM2, TM3, N-terminus, and ECL2 which matches with previous metadynamics binding simulations for cannabinoid receptors. For the proper characterization of intermediate states, the unbinding processes were further extensively sampled using unbiased simulation and umbrella sampling.

Effectiveness of the post processing techniques TRAM and MSM were compared in predicting the ligands with binding affinity and kinetics. MSM predicts the kinetics and thermodynamics from the eigendecomposition of the transition probability matrix. MSM assumes that the local equilibrium is maintained between the Markovian states. However, with limited sampling, this creterion may not valid between local high and low energy states. TRAM tries to solve this issue by combining biased and unbiased simulation, where biased simulations enhances the local sampling to maintain the equilibrium. We observed that with sufficient data both methods performed in similar way in estimating the standard binding free energy. The relative free energy estimated by both methods match with the experimental result within 0.6 kcal/mol. With lesser amount of unbiased data, TRAM predictions of kinetics and thermodynamics remain more consistent than the MSM as the biased simulations help to maintain local equilibrium.

TRAM estimated thermodynamics helped to decipher the differences between the unbinding of NPS MDMB-FUBINACA and classical cannabinoid HU-210. First, for MDMB-FUBINACA, a larger conformational change is observed within the pocket. A metastable intermediate state is observed when the aromatic tail of FUBINACA flip inside the pocket and reorient itself close to the aromatic residues of TM2. It was observed that both linked part and tail part of the ligands can lead the dissociation of the ligand from the receptor. Second, for HU-210, conserved cyclic group leads the dissociation from the receptor. It supports previous simulation where the alkyl side chain of the ligand binds to the receptor first.^17^ Third, aromatic residues in the pocket (F268^ECL2^, F170^2.57^) form major interactions with both HU-210 and MDMB-FUBINACA. Major differences in protein-ligand interactions were observed in TM7. Stronger interactions were observed for the classical cannabinoid HU-210 with TM7, especially polar interaction with S383^7.39^ and hydrophobic interaction with F379^7.35^ compared to MDMB-FUBINACA. This interaction pattern was consistent across other NPS and classical cannabinoids, indicating a universal difference in how these two groups of compounds interact with TM7.

Finally, we demonstrated that the variation in binding pocket interaction leads to the distinct dynamic allosteric communications in the intracellular region. Allosteric communication strength was measured by the variational autoencoder (NRI). NRI network learns the dynamic interactions between residues in the latent space by learning to reconstruct the dynamics. Dynamic allostery measured by the posterior probability of VAE shows that higher allosteric weights from the binding pocket residues to the NPxxY motif region for MDMB-FUBINACA bound CB_1_ increases the probability of triad interaction formation. Since the triad interaction is crucial for *β*-arrestin signaling, these findings align with experimental observations of enhanced *β*-arrestin signaling in NPS-bound receptors. Overall, these data driven computational study helps us to distinguish between the receptor-protein interaction, unbinding mechanism and downstream signaling NPS compared to other classical cannabinoids.

## Methods

### System Preparation

For NPS unbinding simulation, G-protein bound active structure (PDB: 6N4B^45^) was selected as the initial structure. G-protein subunits and Non-Protein residues other than orthosteric ligand MDMB-Fubinaca were removed from the PDB structure file. Missing residues in ICL3 (21 residues, 314-334) and ECL2 (6 residues, 258-263) were modeled sequentially using Remodel protocol of Rosetta loop modeling.^72,73^ In each step, the remodeled structure with least energy was further refined using kinematic closure protocol. ^74^

Starting 108 residues from CB_1_ N-terminus were also missing from the cryo-EM structure. However, it is not feasible to model the entire N-terminus because of two following reasons. ^72^ First, a proper template is not available for modeling N-terminus regions as most of the class A GPCRs do not contain large N-terminus. ^75^ Second, it is challenging to model these large numbers of residues accurately with template-free *ab initio* modeling because of the combinatorial expansions of conformational space. Therefore, the closest 20 residues (89-108) were modeled as membrane proximal regions of the N-terminus were shown to be important for CB_1_ signaling by allosterically modulating ligand affinity. ^76^ Furthermore, Δ89*CB*_1_ (CB_1_ with first 88 residues truncated in N-terminus) have similar ligand binding affinity compared to CB_1_ with full sequence. ^77^ Modeling of this membrane proximal region was also performed Remodel protocol of Rosetta loop modeling. A distance constraint is added during this modeling step between C98^N−term^ and C107^N−term^ to create the disulfide bond between the residues.^76,78^

As the cryo-EM structure of bound MDMB-FUBINACA was known, ligand coordinate of MDMB-FUBINACA was added to the modeled PDB structure. The “Ligand Reader & Modeler” module of CHARMM-GUI was used for ligand (e.g., MDMB-Fubinaca) parameterization using CHARMM General Force Field (CGenFF). ^79^ The ligand bound receptor was embedded in the bilayer membrane and salt solution (extracellular and intracellular region) using CHARMM-GUI.^80^ As CB_1_ is majorly expressed in central nervous system, an average brain membrane composition of asymmetric complex membrane was selected. The membrane composition was obtained from Ingólfsson et al. and proportionally downsized according to our system (Table S1).^81^ 150 mM NaCl salt solution with TIP3P water model was used in the extracellular and intracellular regions. ^82^ CHARMM36m forcefield was used to parameterize the protein, lipid, water, and ions.^83^

For building the classical cannabinoid system, the modeled PDB structure was used. In this case, a classical cannabinoid HU-210 was docked into the orthosteric pocket using Autodock Vina. ^84^ The docked bound pose was selected based on best overalled structure of HU-210 to the experimentally determined crystal structure of another bound classical cannabinoid (Ligand: AM841, PDB:6KPG ^46^) (Figure S13). The classical cannabinoid-bound system was built with identical complex membrane composition, salt concentration, and forcefield with NPS bound system.

Other classical cannabinoids (AMG-41, JWH-133, and O-1317) and NPS (AMB-FUBINACA, CUMYL-FUBINACA, 5F-AMP) bound systems were also set up. These ligands are docked into the orthosteric pocket. Best docking poses were selected based on optimizing the distance between the hydroxyl group of classical cannabinoid (linker oxygen for NPS) to S383^7.39^ and the furtherest tail atom distance to W279^5.43^. These systems also have identical complex membrane composition, salt concentration, and forcefield with previously described systems.

### System minimization and Equilibration

All ligand-bound systems are minimized and equilibrated before the production run. Ten thousand minimization steps were performed with the conjugate gradient method. Six sequential equilibration steps were carried out to stabilize the systems at 300 K temperature and 1 atm pressure for the production simulations. The systems were heated to 300 K in the NVT ensemble in the initial two stages. Each of these steps was performed for 250 ps. Langevin dynamics was used to control the temperature with additional damping and random force. The damping coefficient for the damping force was set as 1/picosecond. The Langevin dynamics was turned off for the hydrogen atoms. All the bonded hydrogen atoms were constrained with the SHAKE algorithm with default parameters of NAMD. ^85^ The integration time step for these two NVT ensemble equilibration was one femtosecond (fs). Harmonic constraints were used for fixing the temperature coupling of the protein residues with constraint scaling term set to 10 in the first NVT ensemble equilibration, followed by Temperature coupling of lipid molecules was also restrained with harmonic force with a force constant equal to 5. CHARMM-GUI Selected Dihedral and Improper bonds were also restrained with an extra bonded term with a force constant of 500 in the first NVT ensemble equilibration, followed by 200. The non-bonded cutoff distance for the van der Waals interactions was set to be 12 Å with a switch distance of 10 Å, at which a switching function is turned on to truncate van der Waals interactions at the cutoff distance. Non-bonded interactions for three consecutively bonded atoms were excluded. The particle mesh Ewald method was implemented for electrostatics calculation with grid size 1 Å.^86^

The next four equilibrations were performed in the NPT ensemble, where pressure was fixed to 1 atm with Langevin piston pressure control. The barostat oscillation period was set to 100 fs with a damping time 50 fs. These four NPT ensemble equilibrations were performed for 250, 500, 500, and 500 ps, respectively. The integration timestep for these equilibration steps was increased to two fs. The constraint for temperature coupling on the protein residues was decreased gradually with the constraint scaling term for the four NPT ensemble simulations set to 2.5, 1.0, 0.5, and 0.1, respectively. Similarly, the restraints on temperature coupling on the lipid molecules were also decreased gradually with force constant for the four NPT ensemble simulations set to 2, 1, 0.2, and 0.0, respectively. Furthermore, the restraints on the CHARMM-GUI Selected dihedral and improper bonds were decreased gradually with force constant for the four steps set to 100, 100, 50, and 0.0, respectively.

### Well-tempered metadynamics

Well-tempered metadynamics was implemented for finding unbinding pathways.^47,87^ Simulations were performed with the Collective variables module (Colvars) of NAMDv2.14. ^88,89^ In metadynamics, a history-dependent biasing potential (*V*_*meta*_(***S***, *t*)) is added to the Hamiltonian of the MD simulation, which discourages the system from revisiting configurations that have already been sampled. ^64,90^ The *V*_*meta*_(***S***, *t*) is a sum of Gaussians deposited along the system trajectory in the CVs space (***S*** = (*S*_1_(*r*), *S*_2_(*r*)..*S*_*d*_(*r*)) as shown in Equation 1, where W, *σ, τ* are Gaussian height, width, and deposition time step, respectively. With a sufficiently long simulation, the bias potential estimates the underlying free energy along the CVs.^64,87^ The Well-tempered metadynamics was introduced to increase the convergence of bias potential by decreasing the Gaussian height with time (Equation 2).^64^ In the Equation 2, *ω* is the bias deposition rate and *ωτ* is equivalent to constant gaussian height for well-tempered metadynamics. The free energy is estimated from the bias potential using Equation 3, where the ΔT is an user defined parameter.

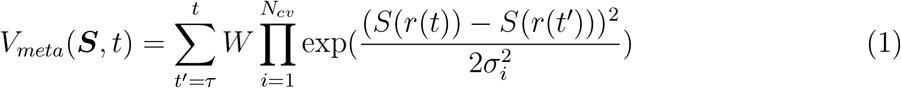

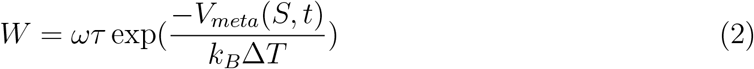

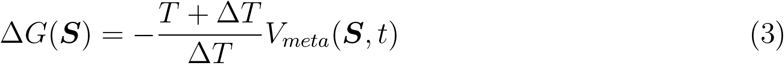

In this work, we selected two collective variables (CVs) for NPS (MDMB-FUBINACA) and classical cannabinoid (HU-210) unbinding according to Mahinthichaichan et al.. ^62^ The first collective variable is the z-direction distance from the converged toggle switch to the center of mass of all heavy carbon atoms of ligands. The second collective variable is the coordination number (CN) as defined in Equation 4 where *d*_*ij*_ represents the distance from ith atom of the ligand and the alpha carbon of jth residue in the binding pocket. Selected binding pocket residues for CN calculation are shown in the Supplementary table. The ΔT, gaussian height (*ωτ*), deposition time step (*τ*) are selected as 4200K, 0.4 kcal/mol and 100 ps, respectively. The gaussian width for the two CVs are set to be 0.5 and 0.1, respectively.

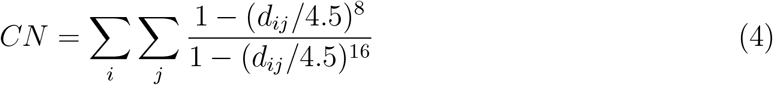

### Umbrella Sampling and Unbiased Sampling

Umbrella sampling was performed along ligand distance from TM5 to capture the unbinding process (Figures S14A and S14B).^55^ The unbinding pathway obtained from the metadynamics was clustered into 300 bins by dividing the selected distances from 5 to 35 Å. The center of each bin was used as the center of each window for umbrella sampling. Five independent structures were selected from each cluster to simulate five independent umbrella runs in each umbrella window. If a cluster does not contain any structure, starting structures for that window were selected from the closest clusters. A constant harmonic biased potential of 10 kcal/mol is used for each window. openMM v7.8 MD engine was used to run the umbrella sampling runs. ^91^ The temperature and pressure of the systems are controlled at 300K and 1 atm by the Langevin thermostats and Monte Carlo barostats. The integration timestep was chosen to be two fs. Movements of the containing Hydrogen atom were constainted using HBonds commands with SHAKE (or SETTLE for water) algorithm. The cutoff distance for Non-bonded interaction other than electrostatic interaction was set to 12 Å, with a switching potential at 10 Å to make the potential to zero smoothly at the cutoff. The particle weld method was used to calculate the long-range electrostatic. Each simulation was run for 20 ns.

Identical starting structures and simulation conditions (Thermostat, barostat, cutoff, electrostatic calculation method, integration timestep, and constraints on Hydrogen bond) were selected for unbiased simulations. openMMv7.8 simulation software was used to run simulations. Each trajectory was run for 100 ns. All the simulations were performed on the distributive computing facility folding@home. ^92^

### Markov State Model

Markov state model (MSM) is used to estimate the thermodynamics and kinetics from the unbiased simulation.^56,93^ MSM characterizes a dynamic process using the transition probability matrix and estimates its relevant thermodynamics and kinetic properties from the eigendecomposition of this matrix. This matrix is usually calculated using either maximum likelihood or Bayesian approach. ^56,94^ The prevalence of MSM as a post-processing technique for MD simulations was due to its reliance on only local equilibration of MD trajectories to predict the global equilibrium properties. ^95,96^ Hence, MSM can combine information from distinct short trajectories, which can only attain the local equilibrium. ^97–99^

The following steps are taken for the practical implementation of the MSM from the MD data.^4,17,100–102^

1. Each frame obtained from the MD simulation was featurized using features important for capturing the conformational ensemble. In this case, the unbinding process for each ligand was featurized using distances that characterize the ligand distances to the binding pocket and binding pocket conformational change. Specifically, all heavy atom distances from each the C*α* carbon atom of all binding pocket residues were calculated (Figure S15). Additionally, all possible combinations of C*α* carbon atom distances between all the binding pocket residues were included to capture the binding pocket motion. Feature calculations were performed with the python library MDTraj v1.9.8.^103^ The total number features selected for MSM building of MDMB-FUBINACA and HU-210 are 297 and 288, respectively.
2. Dimensionality reduction was performed using time-independent component analysis (TICA).^104,105^ We found the orthogonal projections (time-independent components) with TICA, which are linear combinations of the slowest features. In tIC space, two spatially close points are kinetically close. The lagtime selected for tiC building was 5ns.
3. Clustering was performed on the tICs using k-means clustering algorithms to discretize the space into Markovian states.
4. Lag time for the MSM was calculated by estimating the shortest time at which the timescale of the slowest processes has converged to a particular value (Figures S16A and S16B).
5. To optimize MSM based on the cluster numbers and tIC components on which clustering is performed, we calculated the VAMP-2 score from the MSM, where VAMP stands for Variational Approach for Markov Processes (Figure S17). ^106^ For a reversible MSM, this score represents the summation of the square of the k slowest eigenvalues, where k is a hyperparameter. Closer the eigenvalue to 1, the corresponding eigenvector captures a slower process. Therefore, we optimize the MSM by maximizing the VAMP-2 score.
6. To validate the Markovian property of our optimized models, Chapman–Kolmogorov test (c-k) test was performed (Figure S18). C-K test states that for a Markov model kth power of *P* (*τ*) needs to be equal transition probability matrix determined at *kτ* time (*P* (*τ*)^*k*^ ≈ *P* (*kτ*)). We showed that differences between the elements of transition probability matrix at higher lag times remain relatively small.

Dimensionality reduction, clustering Markov state model building, and VAMP-2 calculations are performed with the pyEMMA v2.5.6 library.^107^ The optimized MSM for MDMB-FUBINACA unbinding simulations were built with 700 clusters, 7 tiCs and 35 ns of lag time. For HU-210, optimized MSM were build with 800 clusters, 6 tiCs and 35 ns of lag time.

### Transition-based reweighting analysis method

Markov State Models have been extensively used to investigate the protein-ligand binding process.^17,67,108–113^ However, these studies were mainly performed for ligands with high off-rates which could be sampled using the unbiased trajectories. For ligand with low off rates, the use of reversible transition matrix would yield incorrect estimates of unbinding kinetics. Therefore, we use The Transition-based reweighting analysis (TRAM)^48,114^ method to accurately estimate the unbinding kinetics of new psychoactive substances. TRAM is a thermodynamics and kinetics estimator method, which, unlike MSM, can combine unbiased and biased simulation data to estimate thermodynamics and kinetics. TRAM utilizes the advantages of the local equilibrium approximation of MSM and the benefits of biased simulations to enforce local equilibrium in interstate transitions where it is difficult to attain.

As the simulations are obtained from multiple ensembles (biased and unbiased), it is paramount to classify the MD frames (or the conformations) based on which ensemble it belongs to. Each ensemble represents simulations that are performed with identical Hamiltonian energy functions. Therefore, unbiased simulations are considered as one ensemble, whereas, in umbrella sampling, each biasing window is considered a single ensemble.

Like MSM, in TRAM, the conformational space is also discretized into non-overlapping states. The interstate transitions should follow the following relationship shown in the Equation 5, where 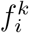 is the local free energy of the ith state and kth ensemble. The term 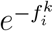 is proportional to the stationary density (*µ*(*x*)) of state *S*_*i*_ in ensemble *k*. The *µ*(*x*) of each conformation (*x*) of *S*_*i*_ is weighted with negative exponential of bias energy (*b*^*k*^(*x*)) on x in ensemble 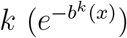 (Equation 6).

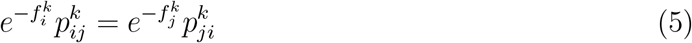

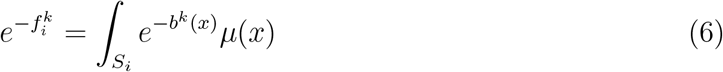

To obtain kinetics and thermodynamics information from TRAM, we have to derive interstate transitions 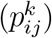 and the stationary density of the entire ensemble (*µ*(*x*)), where both the terms follow normalization constraint (Equation 7 and 8). Therefore, there are *m*^2^*K* + *X* unknown variables. Therefore, to solve these unknown variables, the maximum likelihood approach has been considered, where the likelihood function is defined as Equation 9, which is the combination of the likelihood function of MSM and local equilibrium. This maximum likelihood problem was subjected to the constraints of Equation 5, 7 and 8.

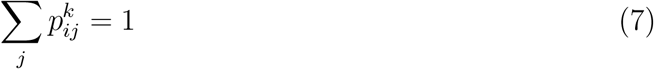

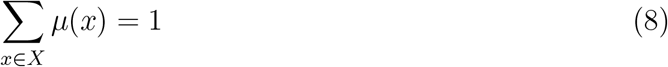

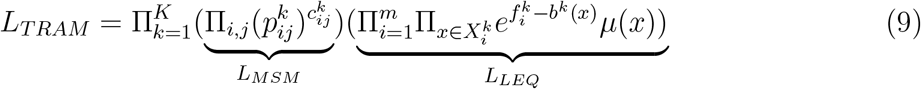

Wu et al. showed that the solution of these maximum-likelihood problem can be turned into system of non-linear algebraic equations (Equation 10, 11 and 12), where 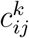 is count of the interstate transitions between state *S*_*i*_ and *S*_*j*_ in ensemble *k*. This system of equations are solved iteratively to estimate 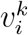 and 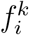, which provides the prediction of 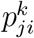 and *µ*(*x*) (Equation 13 and 14).

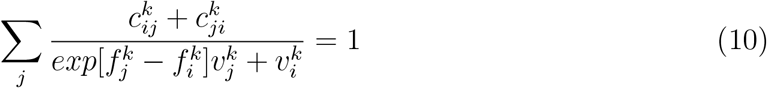

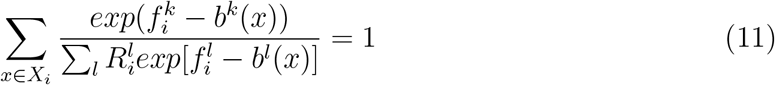

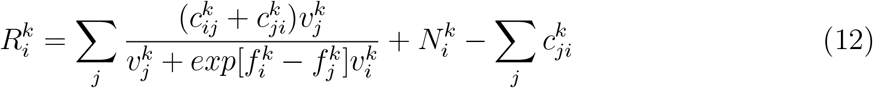

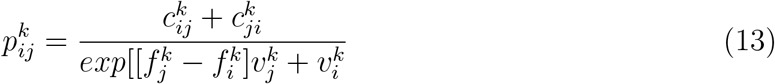

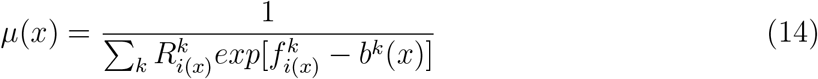

We used the python package pyEMMA v2.5.6 for the practical implementation of TRAM.^107^ For calculating transition counts in ensemble 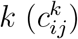, the lagtime of 15 ns was chosen. In the implementation, we need to preprocess each trajectory into three arrays.

1. One of the arrays represents the spatial discretization of each trajectory frame, where each frame belongs to a particular state. Therefore, each element can take values from 0 to m-1 (m is the cluster number). Before the discretization of the space, time-independent component analysis was performed on the biased and unbiased data separately. The number of tIC components for each system was selected based on the number of the tIC components of optimized MSM. Each frame from the unbiased simulation is represented by the unbiased tICs, concatenated with its feature projections on the biased tICs. Similarly, each frame from the biased simulation is represented by its feature projections on the unbiased tic, concatenated with biased tic projection. Therefore, NPS unbinding simulations have 14 tICs, whereas classical cannabinoid unbinding simulations have 12 tICs. The number of clusters is also obtained from the optimized MSMs.
2. Another array represents the corresponding ensemble to which each trajectory frame belongs. There are 300 windows for the umbrella sampling. Therefore, there are 301 ensembles, as the unbiased simulations represent a separate ensemble.
3. Third array represents the corresponding bias potential (*b*_*k*_(*x*)) a particular frame feels if it were to be in a particular ensemble. For umbrella sampling, the biased potential is represented as Equation 15, where *c*_*k*_ is selected to be 10 kcal/mol and *y*_*k*_ is the center of each umbrella window.

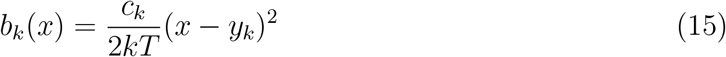

### Transition Path Theory

Transition path theory (TPT) analysis is applied to calculate the transition pathway and timescale between different macrostates, representing different configurational spaces in the unbinding process.^115,116^ In this work, we define macrostates as a collection of Markovian states present in the area of interest in the unbinding free energy landscape. An essential concept of transition path theory is the committer probability 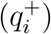, which is defined as the probability of any Markovian state reaching the final metastable state before it returns to the initial state. Therefore, the Markovian states present in metastable state B has a committer probability of 1. It has been shown that committer probability follows the following system of linear equation as shown in Equation 16, where *P*_*ik*_ is the transition probability between state *S*_*i*_ and *S*_*j*_ as discussed in the previous section.

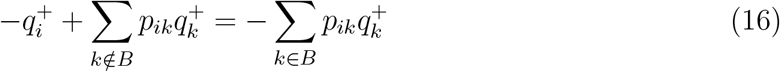

In this work, the quantity of interest from TPT is the timescale (or rate) between the metastable state transitions as shown in Equation 17, where *π*_*i*_ is the stationary probability of state *S*_*i*_. TPT calculations were performed by PyEMMA v2.5.6. ^107^

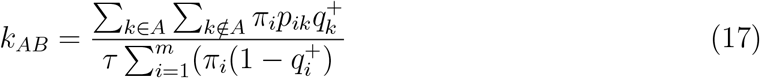

### K-L Divergence Analysis

Kullback–Leibler divergence (K-L divergence) analysis was performed to show the structural differences in protein conformations in different macrostates^4,117^. In this study, this technique was used to calculate the difference in the pairwise inverse distance distributions between macrostates. Each macrostate was represented by 1000 frames that were selected proportional to their TRAM weighted probabilities. Although K-L divergence is an asymmetric measurement, for this study, we used a symmetric version of the K-L divergence by taking the average between two macrostates. Per residue contribution of K-L divergence was calculated by taking the sum of all the pairwise distances corresponding to that residue. This analysis was performed by in-house Python code.

### Trajectory Analysis

Python package GetContacts is used to perform the contact calculation. ^118^ Linear interaction energy analysis was performed to calculate the interaction energy between ligand and receptor using AMBERTools CPPTraj v18.01.^119,120^ Trajectory visualization and figure generation are performed with VMD v1.9.3. ^121^

### Deep Learning Network for Allosteric Prediction

Neural relational inference (NRI) network was implemented to predict allosteric dependence between the residues in the different parts of the receptors.^57,122^ This network is a Variational autoencoder (VAE) comprising encoding and decoding parts. ^123^ The encoder (*q*_*ϕ*_(***z***|***x***)) takes the input C*α* coordinates of protein conformations at time t (***x***_***t***_) and tries to learn the interactions between two residues (*z*_*ij*_) in the protein as a latent space. The decoder (*p*_*θ*_(***x***|***z***)) network try to regenerate the protein conformation at time *t* + *τ* (***x***_***t*+*τ***_). Similar to other VAE, the learning process maximizes the evidence lower bound (ELBO) as shown in Equation 18, where *p*_*θ*_(***z***) represents the prior distribution for ***z***. Here, the prior distribution is selected as default presented in the original paper, where it is represented as a categorical distribution with *K* = 4 (*P*_1_ = 0.91, *P*_2_ = 0.03, *P*_3_ = 0.03, *P*_4_ = 0.03).

As shown in Equation, the ELBO consists of two terms. In the first term, further mathematical derivations can show that the first term can be represented as the the reconstruction error (Equation 19), where *σ*^2^ is variance of the distribution, a user defined parameter.

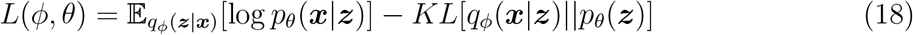

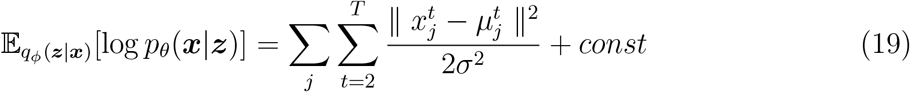

The second term is also called regularization term which is the the K-L divergence between estimated posterior (*q*_*ϕ*_(***z***|***x***)) and prior distribution (*p*_*θ*_(***z***)) (Equation 20). As the prior distribution is a categorical distribution, the K-L divergence becomes entropy of the posterior distribution. We obtained the code for the NRI network from the GitHub implementation and kept most of hyperparameters as default for our training, except for decreasing the hidden layer size to 64.^124^ From each unbinding simulations, 10 unbiased trajectories were selected where the ligand remain in the bound pose. Each trajectory has a length of 100 ns. Both cases, the *τ* was selected to be 5 ns. The allosteric weights (posterior probability) were obtained from the validation data (2 trajectories), where training was performed with remaining 8 trajectories (Figure S20). This procedure was repeated three times, where training and validation data were selected randomly.

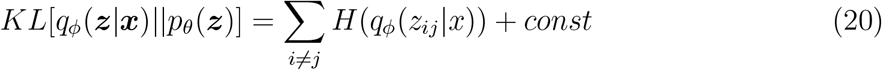

### Mutual Information Estimation

Mutual information between dynamics of residue pairs was computed based on the backbone dihedral angles, as this provides a metric that is independent of the relative distances between residues. The calculations were done on same trajectory data as NRI analysis. Python package MDEntropy was used for estimating mutual information between backbone dihedral angles of two residues.^125^

### Standard Binding Free Energy Calculations

To calculate the standard binding free energy from simulation, we adopted a procedure described in Buch et al.. ^67^ In this procedure, a volumetric correction term is added to the PMF to calculate the final binding free energy (Equation 21). The volumetric correction term is used to predict the free energy at the standard condition (1M) as shown in Equation 22, where *V*_*o*_ corresponds to the volume of a molecule should occupy at the standard condition (1661 *Å*^3^) and *V*_*u*_ is the volume of the unbounded state in the simulation box. The expression for the PMF contribution of the free energy is shown is in Equation 23, where the denominator of the equation can be represented as 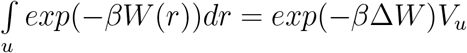. Therefore, the final derivation of Δ*G* is shown in Equation 24

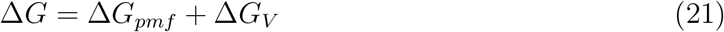

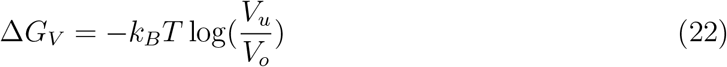

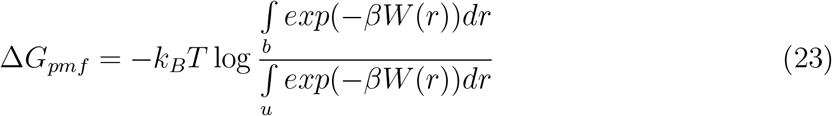

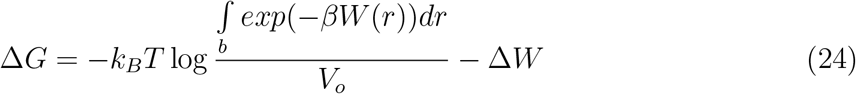

In this work, to estimate the free energy (Δ*G*) x, y, z component of the ligand center of mass is calculated compared to the center of mass of the alpha carbons of binding pocket residues. The three dimensional space was descretized into 25 × 25 × 50 bins and each bin is weighted using TRAM calculated probability density. Depth of the pmf (Δ*W*) was calculated by averaging the pmf of the 100 bins with highest pmf in the bulk. To evaluate the weighted binding volume 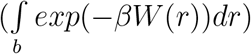, we selected the bins with pmf less than 1 kcal/mol.

## Acknowledgements

D.S. acknowledges support from NIGMS MIRA award R35GM-142745 and NSF Early CAREER Award (MCB-1845606). S.D. and D.S. thank folding@home donors for providing computational resources for the study.

## Code and data availability

Unbinding simulation trajectories and topology files that have been used for the analysis can be obtained from https://doi.org/10.5061/dryad.4f4qrfjq5. Python scripts and necessary files to generate the figures is provided in the github repository. https://github.com/ShuklaGroup/Dutta_Shukla_Cannabinoid_2023a.git

