## Supporting Information for "Characterization of binding kinetics and intracellular signaling of new psychoactive substances targeting cannabinoid receptor using transition-based reweighting method"

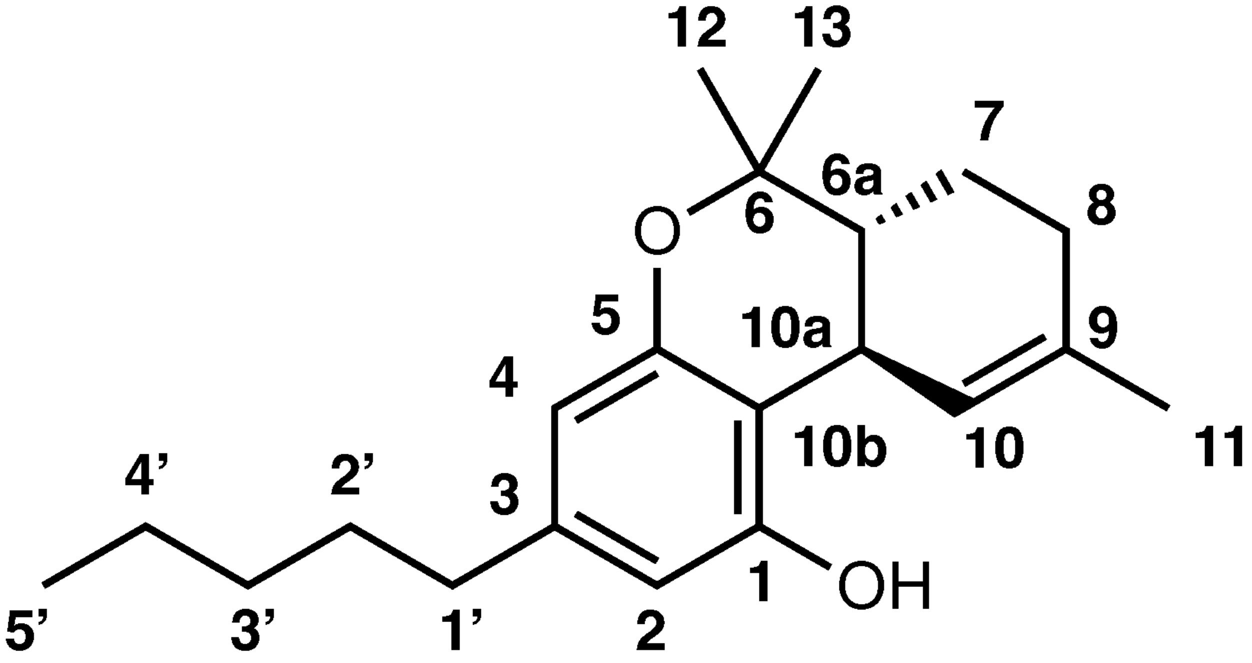


Figure S1: Atom numbering scheme of classical cannabinoid. ($\Delta^9$-Tetrahydrocannabinol)

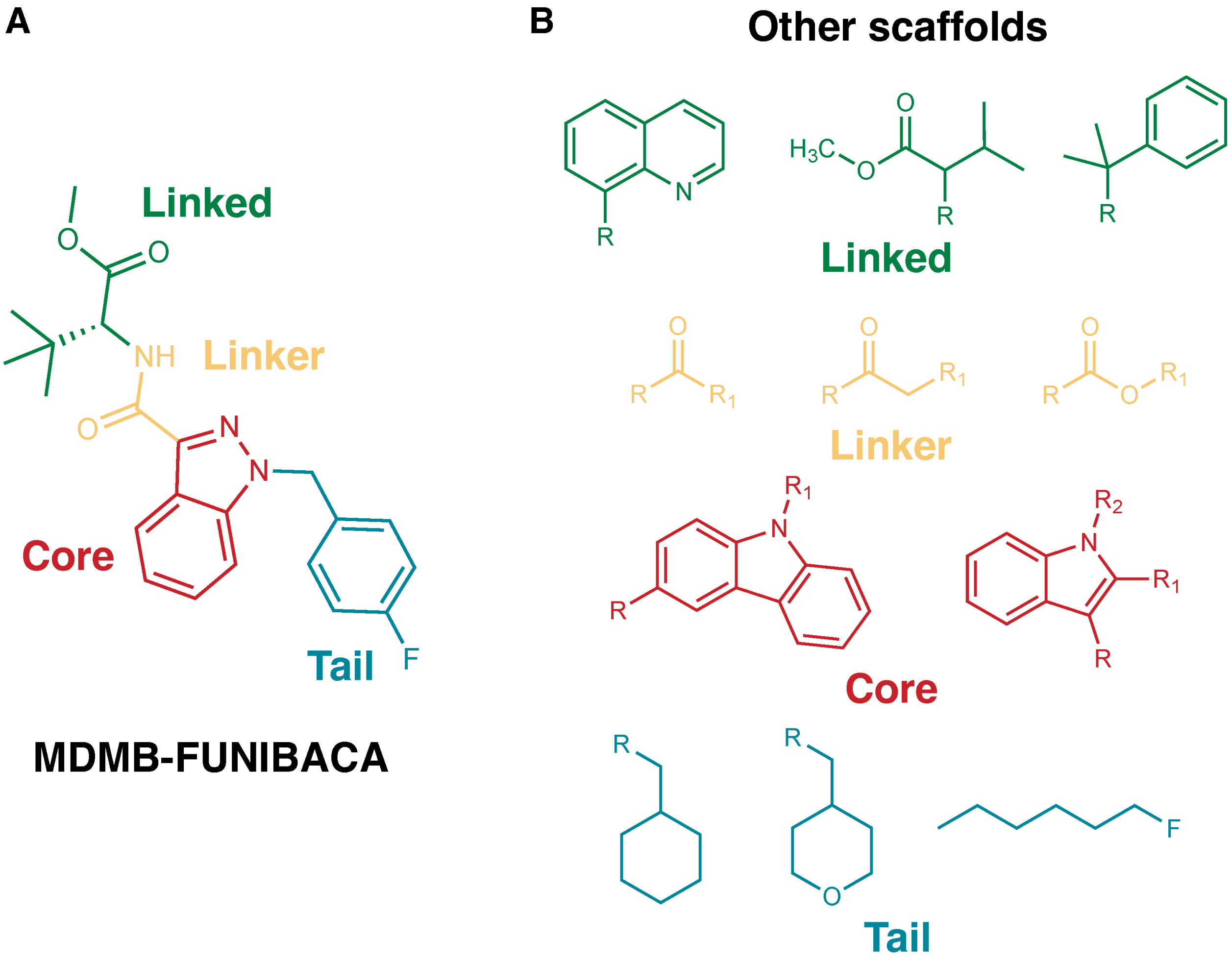


Figure S2: Four pharmacophore components (Linked: Green, Linker: Orange, Core: Red, Tail: Cyan) of NPS synthetic cannabinoid MDMB-FUBINACA are shown in different color (A). Existing common scaffolds of other NPS in four pharmacophore components (B)

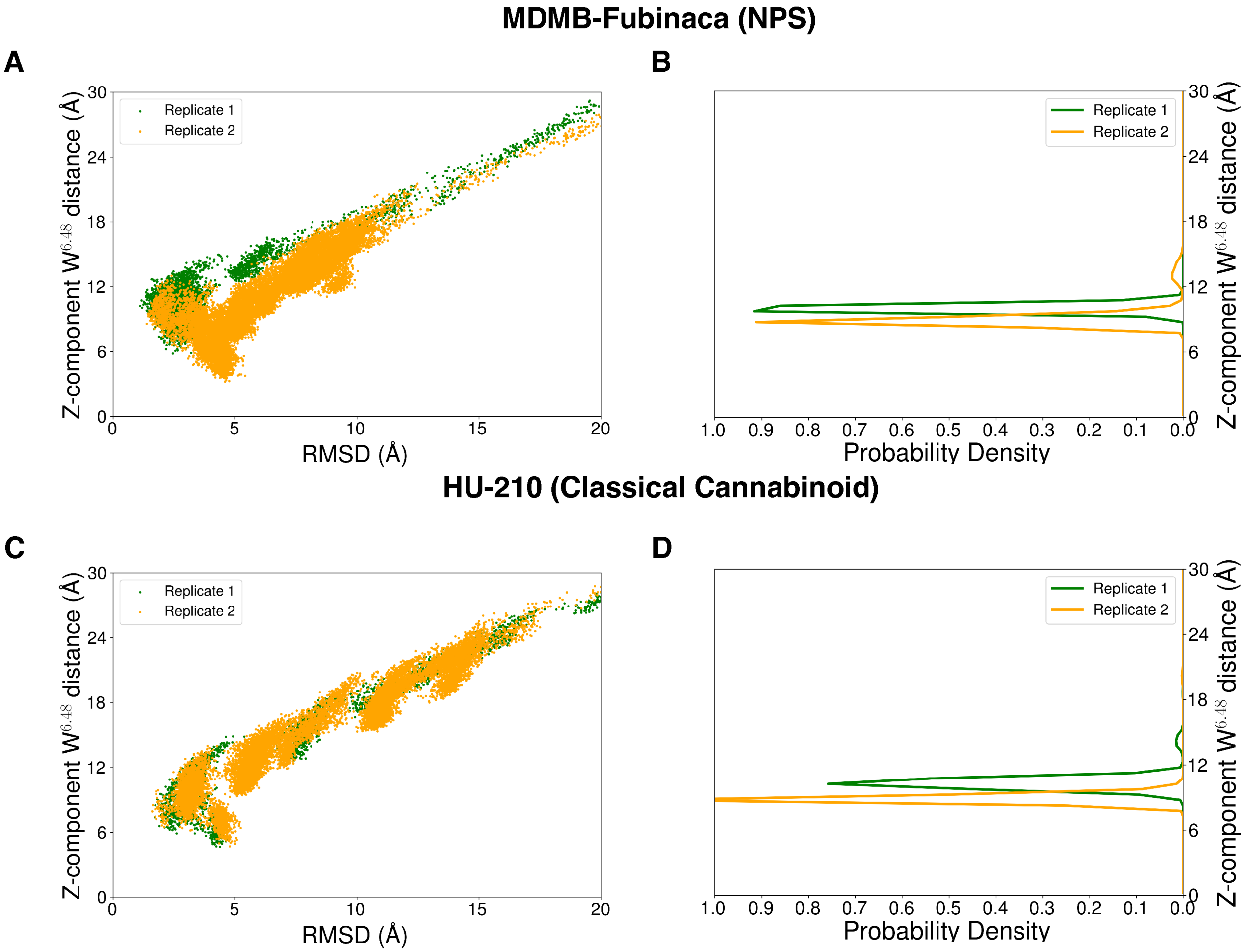


Figure S3: Unbinding ensemble of the MDMB-FUBINACA (A) and HU-210 (C) are projected as 2-D scatter plots where Z-component distance of ligand center of mass from W356$^{6.48}$ is plotted against the ligand RMSD. These ensembles were obtained by running well-tempered metadynamics. Reweighted probability densities are plotted with respect to the Z-component distance of ligand center of mass from W356$^{6.48}$ for MDMB-FUBINACA (B) and HU-210 (D). Measured qualities from two simulation replicas for each system are shown in different colors on the same plot.

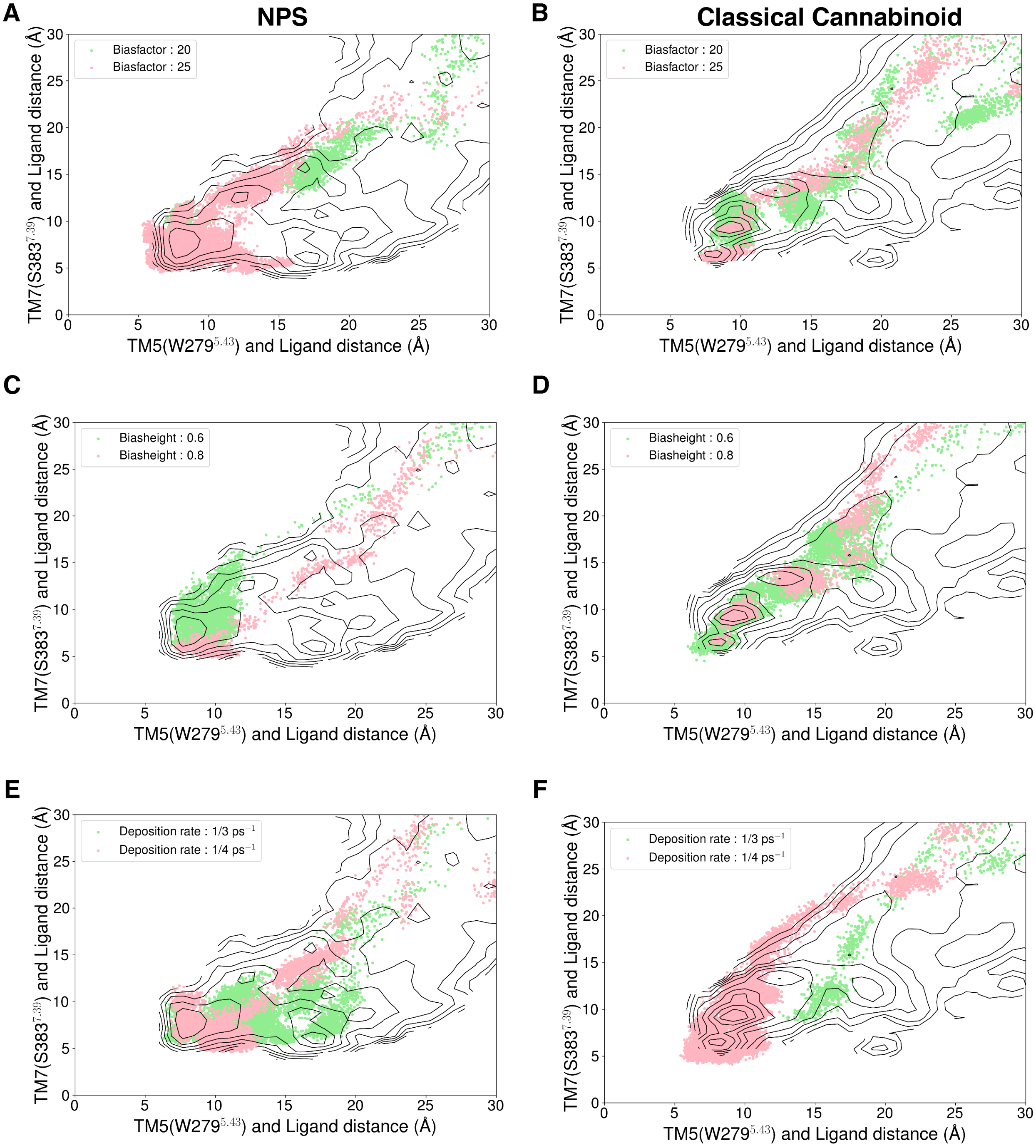


Figure S4: Well-tempered metadynamics runs with different parameters were projected as a scatter plot on top of TRAM weighted two-dimensional projection of unbinding free energy landscape for MDMB-FUBINACA (A, C, E) and HU-210 (B, D, F). For MDMB-FUBINACA, distance between TM5 (W279$^{5.43}$-C$\alpha$) and tail part of the ligand is plotted against the distance between TM7 (S383$^{7.39}$-C$\alpha$) and ligand linked part. For HU-210, distance between the TM5 (W279$^{5.43}$-C$\alpha$) and tail is plotted against the TM7 (S383$^{7.39}$-C$\alpha$) and cyclohexenyl ring of the ligand.

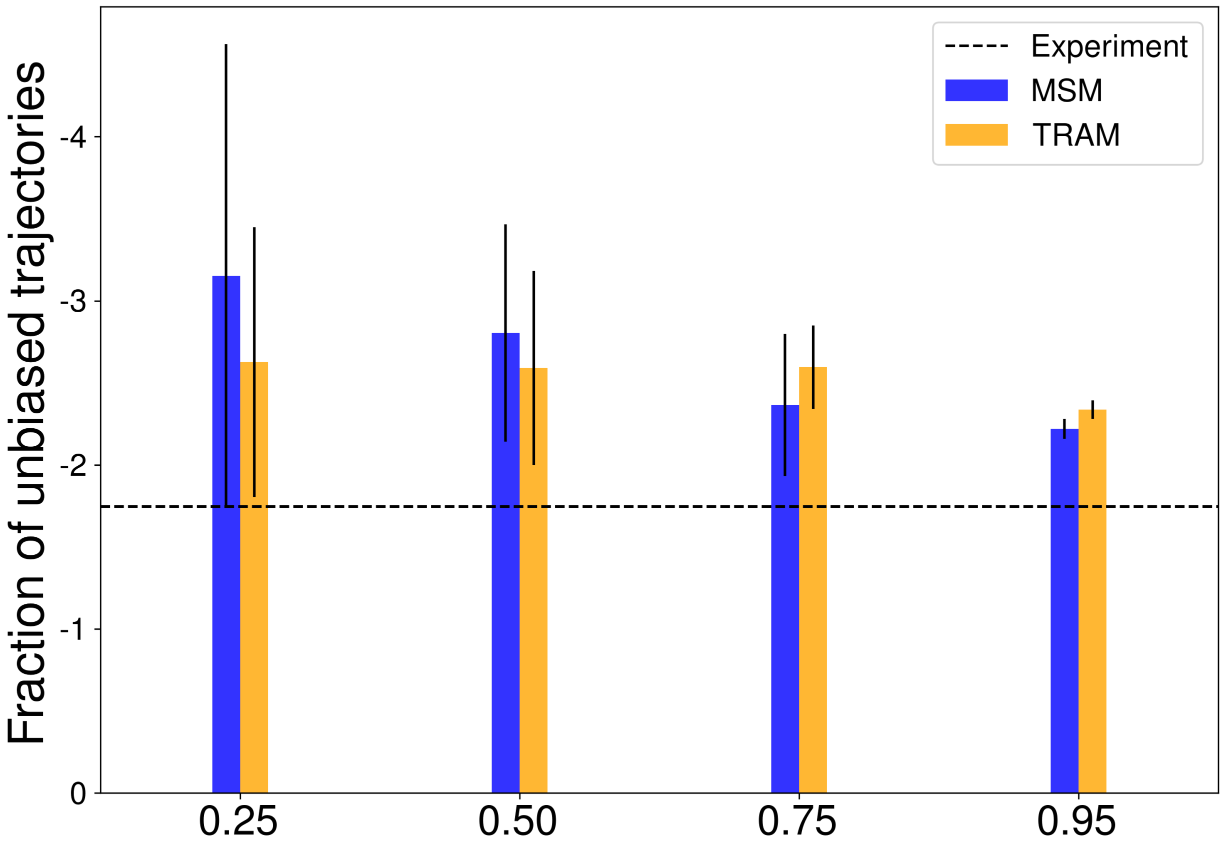


Figure S5: Difference in the $\Delta\Delta G$ of the two ligands is plotted as box plot against fraction of unbiased trajectories used for the estimations. MSM and TRAM estimations are shown using blue and orange colors, respectively.

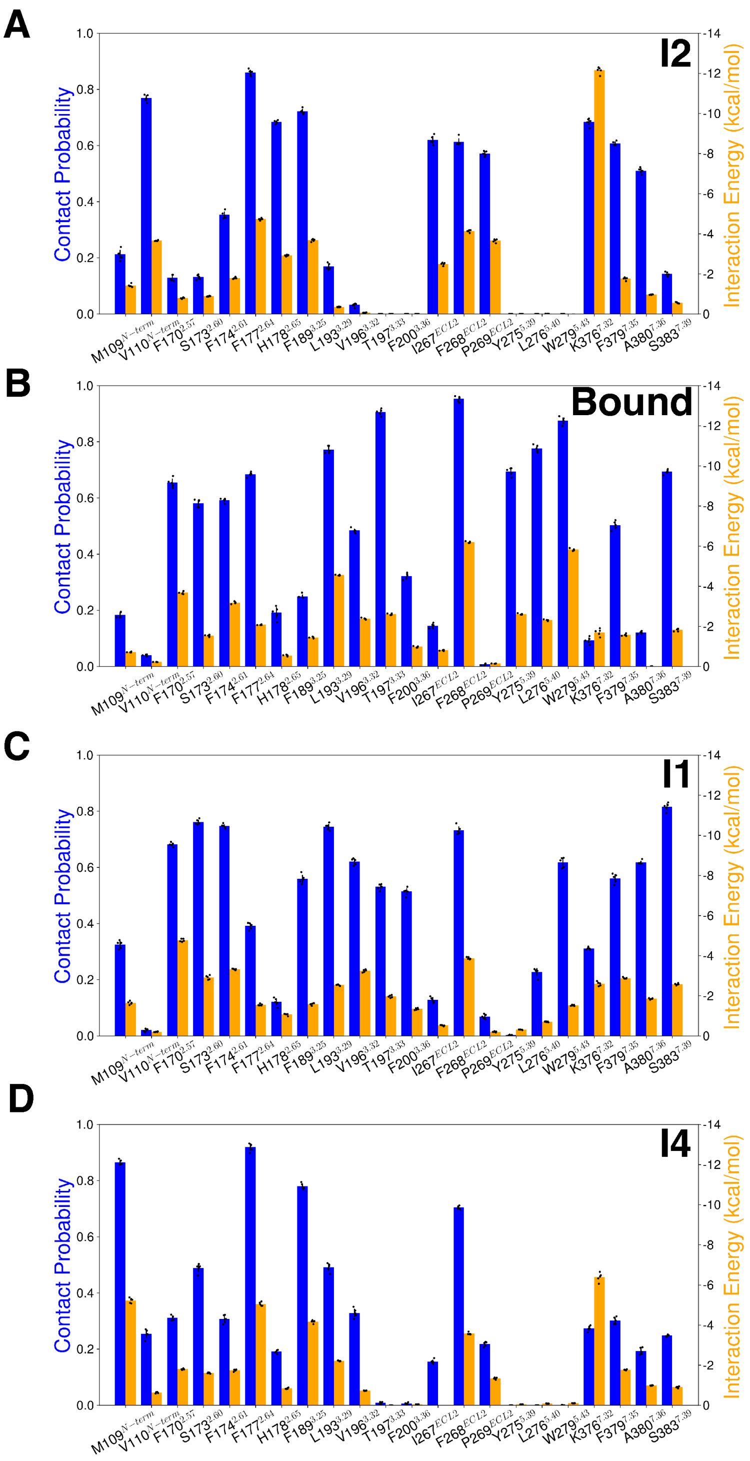


Figure S6: The contact probabilities with binding pocket residues and corresponding interaction energies of MDMB-FUBINACA are shown for different macrostates, where ligand maintains contact with the receptor. The macrostates presented here are in the order of I2 (A), Bound (B), I1(C), and I3(D) to clearly distinguish the two unbinding mechanisms. For each macrostate, ligand and binding pocket residue interaction probabilities (color: blue) and energies (color: orange) are plotted as bar plots.

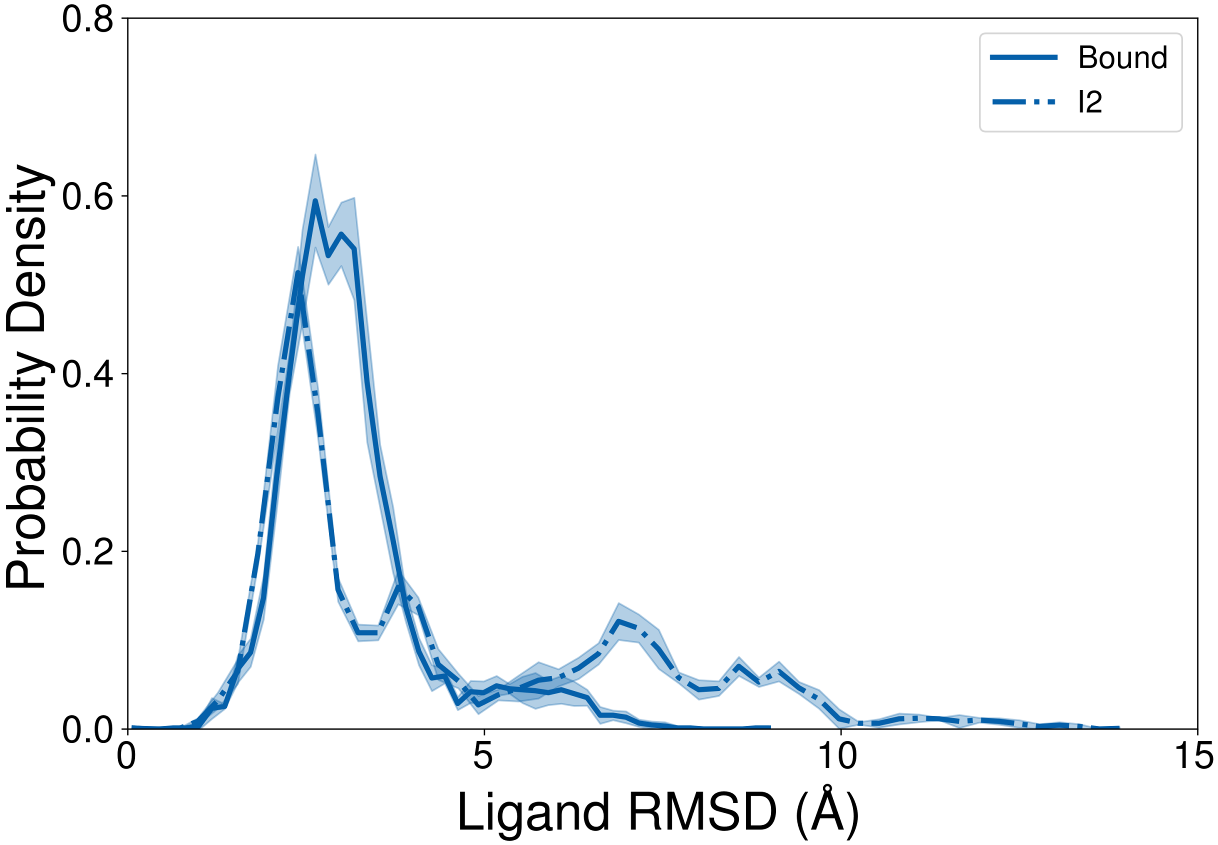


Figure S7: Root mean square deviation of the MDMB-FUBINACA was shown as a density plot for Bound and I2 macrostates. Errors are calculated from 5 bootstrapped samples, where each sample contains 1000 conformations representing the macrostate. These 1000 conformations are selected based on the probability density of the microstates belonging to the macrostate.

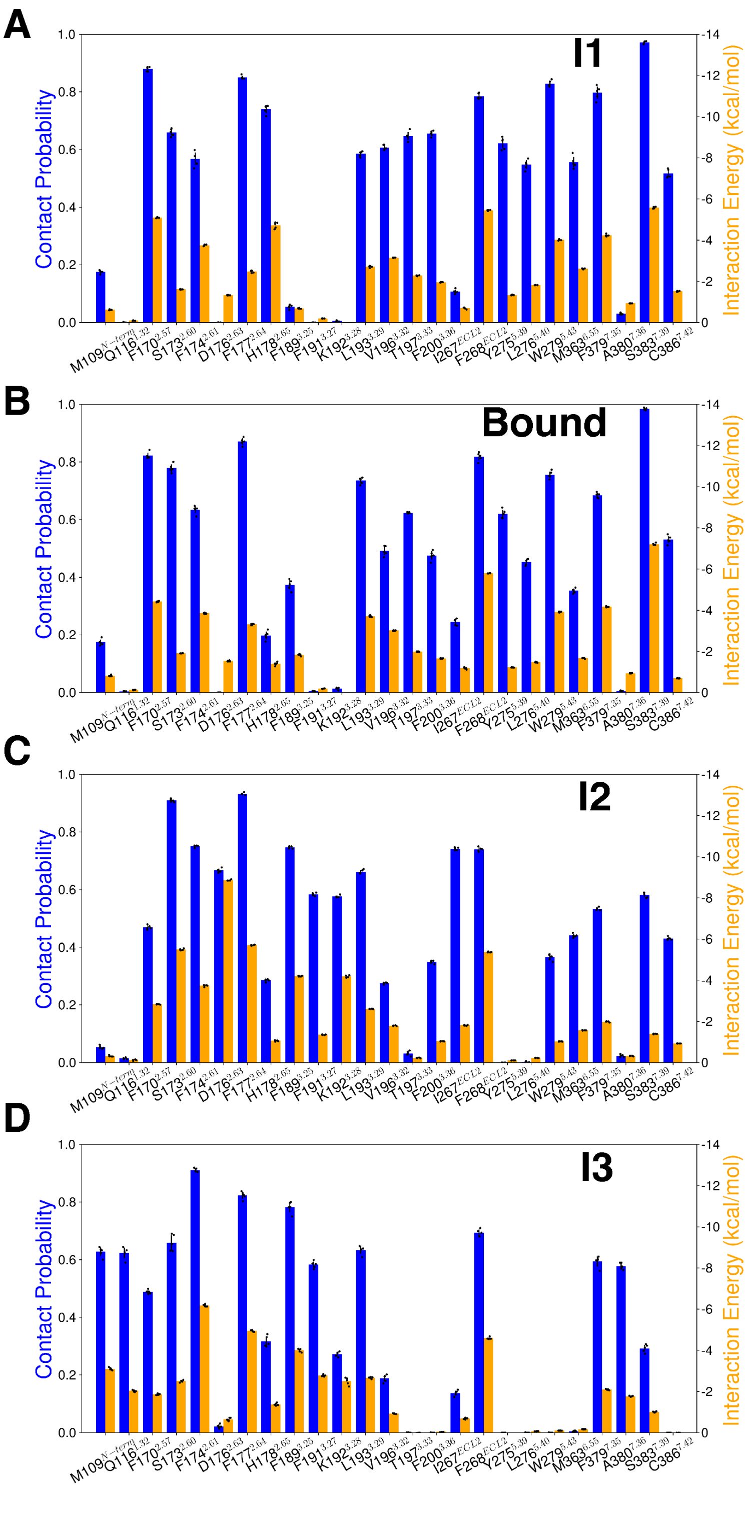


Figure S8: The contact probabilities with binding pocket residues and corresponding interaction energies of HU-210 are shown for different macrostates, where ligand maintains contact with the receptor. The macrostates presented here are in the order of I1 (A), Bound (B), I2(C), and I3(D) to clearly distinguish the two unbinding mechanisms. For each macrostate, ligand and binding pocket residue interaction probabilities (color: blue) and energies (color: orange) are plotted as bar plots.

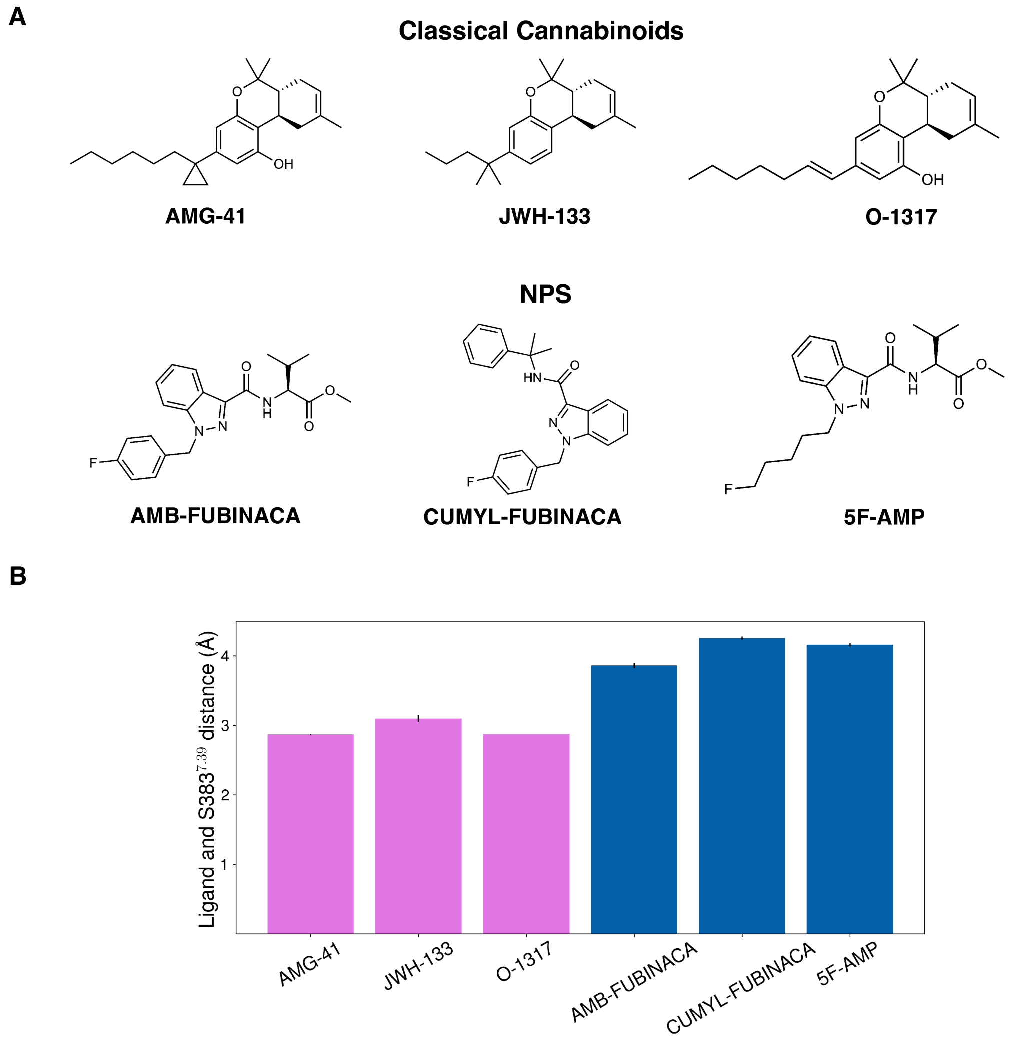


Figure S9: (A) Classical cannabinoids and NPS molecules that have been used to perform unbiased ligand-bound simulations. (B) Equivalent polar interaction distances for NPS and classical cannabinoids are shown as bar plot. For each NPS, the distance from S383$^{7.39}$(O$\gamma$) is calculated from the linker oxygen atom. For each classical cannabinoid the distance from S383$^{7.39}$(O$\gamma$) is calculated from hydroxyl oxygen (or equivalent hydrogen) bound to C1 carbon. The error bar is calculated using bootstrapping approach with 80% of all trajectories. 3 bootstrapped samples are used for the error calculations.

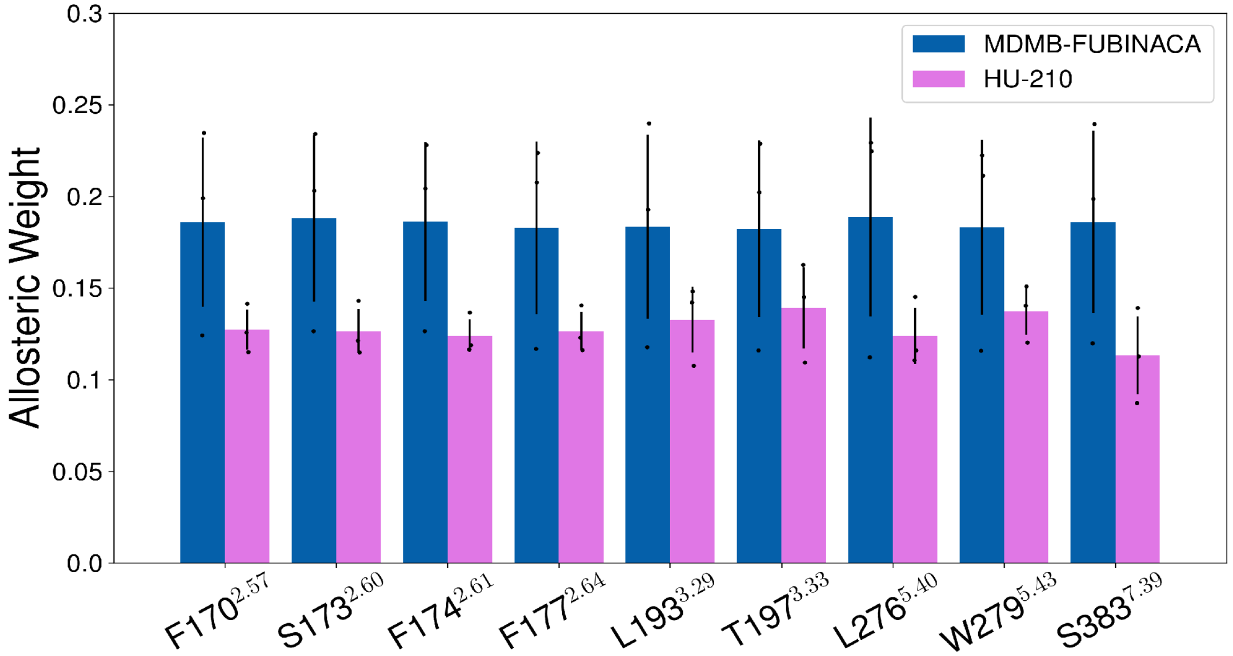


Figure S10: Allosteric weights between the binding pocket residues and the NPxxY motif are plotted as bar plots for HU-210 (color: purple) and MDMB-FUBINACA (color: blue) bound trajectories. Allosteric weights are estimated from the posterior probability of the NRI network. Error in the allosteric weights is calculated by training the network 3 different training data.

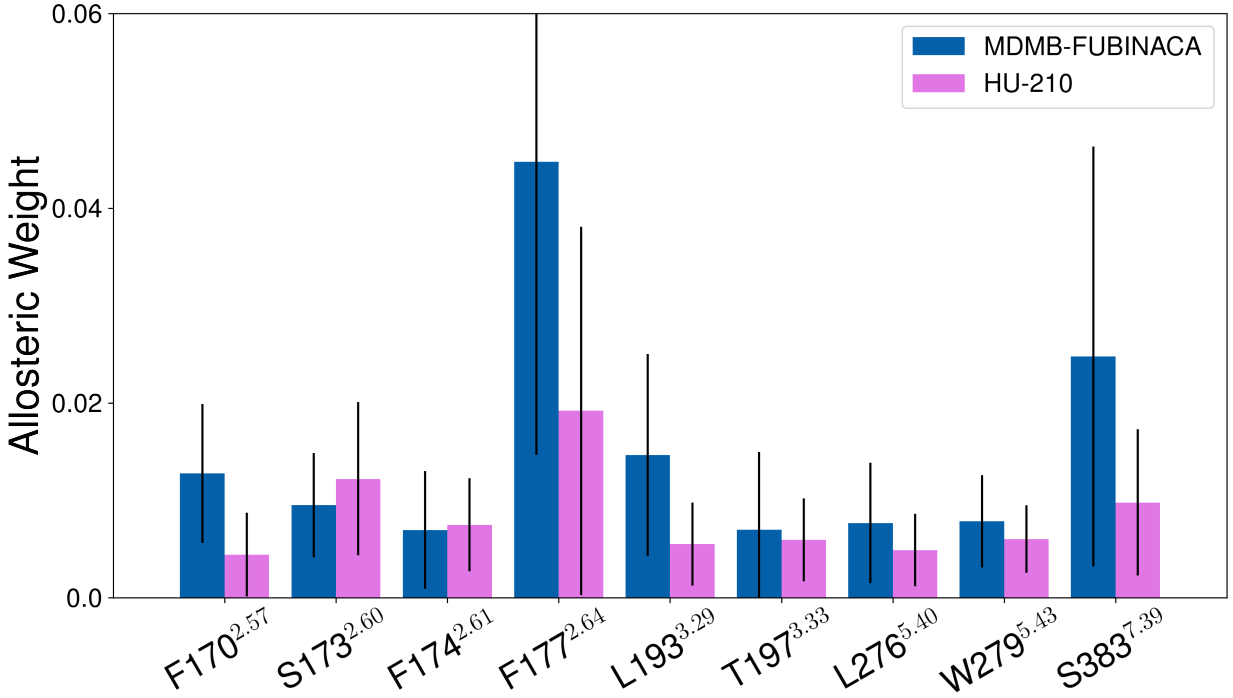


Figure S11: Allosteric weights between the binding pocket residues and the NPxxY motif are plotted as bar plots for HU-210 (color: purple) and MDMB-FUBINACA (color: blue) bound trajectories. These weights were calculated using

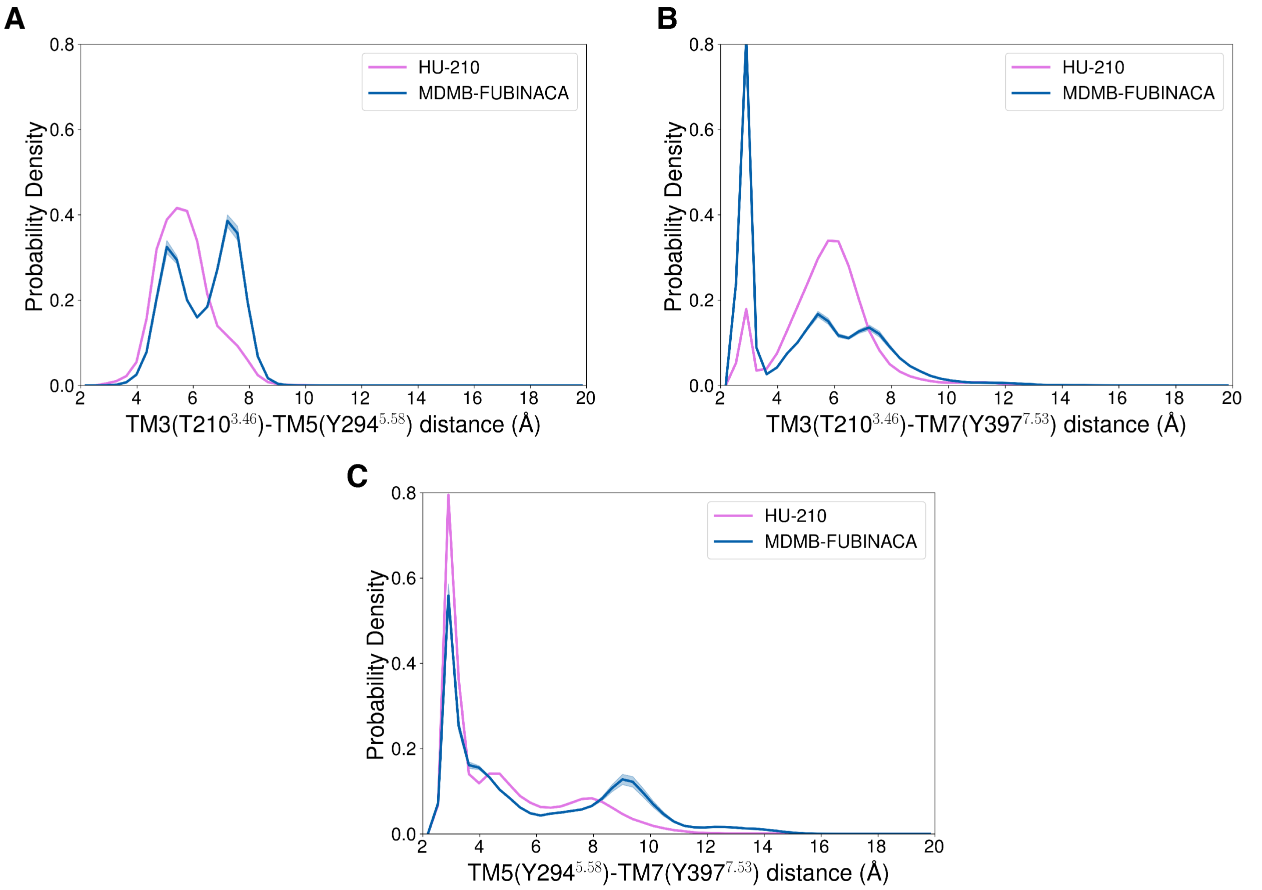


Figure S12: TRAM weighted probability densities of pairwise distances between the residues involved in the triad interaction (Y397$^{7.53}$, Y294$^{5.58}$, T210$^{3.46}$) are plotted for HU-210 (color: purple) and MDMB-FUBINACA (color: blue) unbinding ensemble.

Table S1: Assymetric average brain membrane composition used in MD simulation

| Head Group | Lipid Group | Upper Leaflet | Lower Leaflet | Total |
|---|---|---|---|---|
| phosphatidylcholine | DPPC | 8 | 5 | 13 |
| | POPC | 14 | 7 | 21 |
| | DOPC | 4 | 2 | 6 |
| | PAPC | 7 | 4 | 11 |
| | PDoPC | 1 | 0 | 1 |
| phosphatidylethanolamine | POPE | 2 | 4 | 6 |
| | PAPE | 5 | 9 | 14 |
| | PDoPE | 8 | 15 | 23 |
| sphingolipid | SSM | 9 | 3 | 12 |
| | OSM | 1 | 0 | 1 |
| | NSM | 2 | 0 | 2 |
| phosphatidylserine | DPPS | 0 | 1 | 1 |
| | POPS | 0 | 6 | 6 |
| | PAPS | 0 | 6 | 6 |
| Glycolipid | GM1 | 2 | 0 | 2 |
| | GM3 | 2 | 0 | 2 |
| phosphatidylinositol | POPI | 0 | 5 | 5 |
| | PIPI | 0 | 2 | 2 |
| Ceramide | CER180 | 1 | 1 | 2 |
| Sterol | Cholesterol | 62 | 58 | 120 |
| Total | | 128 | 128 | 256 |

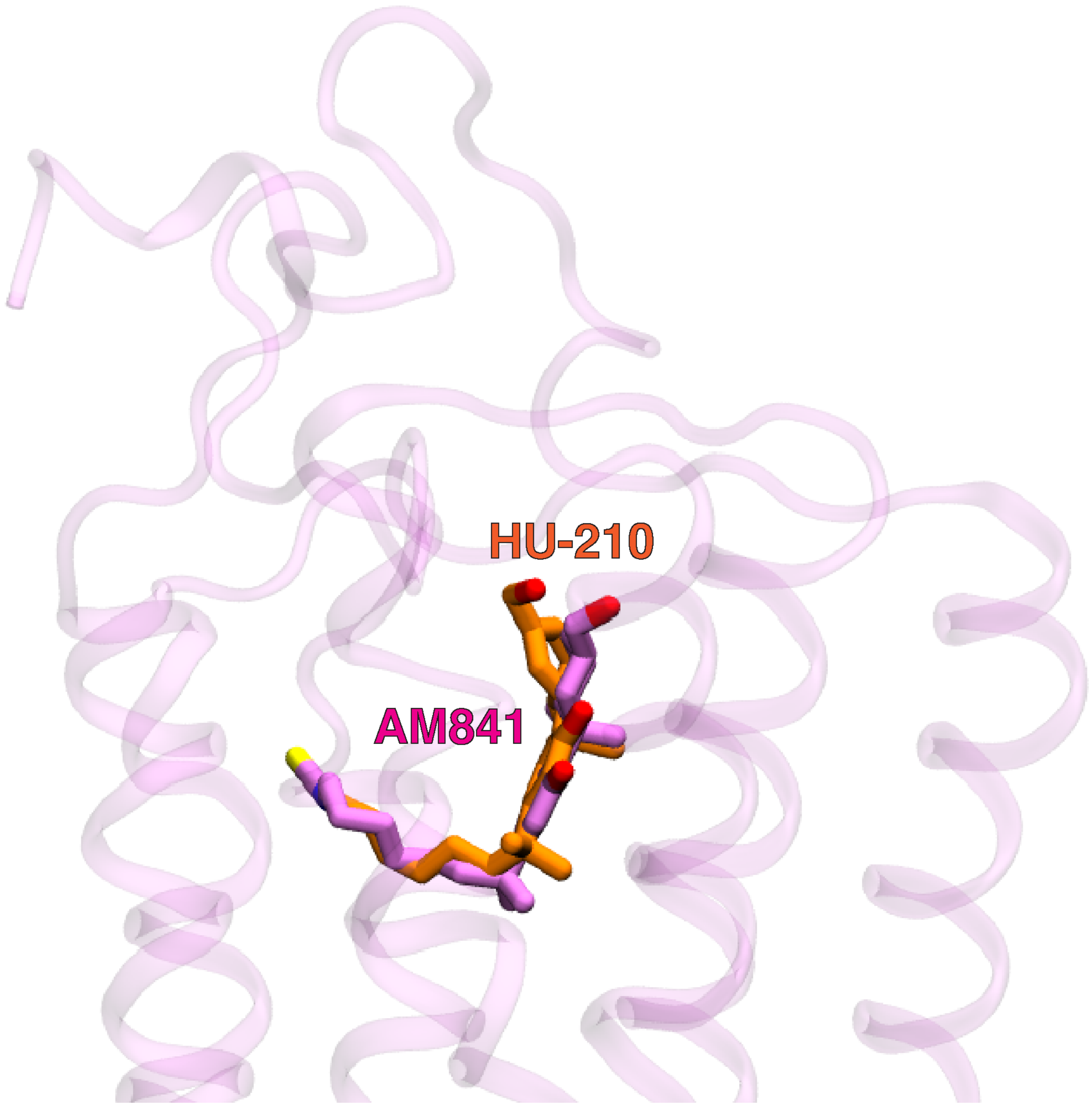


Figure S13: Structural superposition of HU-210 (color: Orange) docked $CB_1$ with active $CB_1$ cryo-EM structure (PDB ID: 6KPG, ligand: AM841). Protein is shown as cartoon (color: purple). Ligands are shown as sticks.

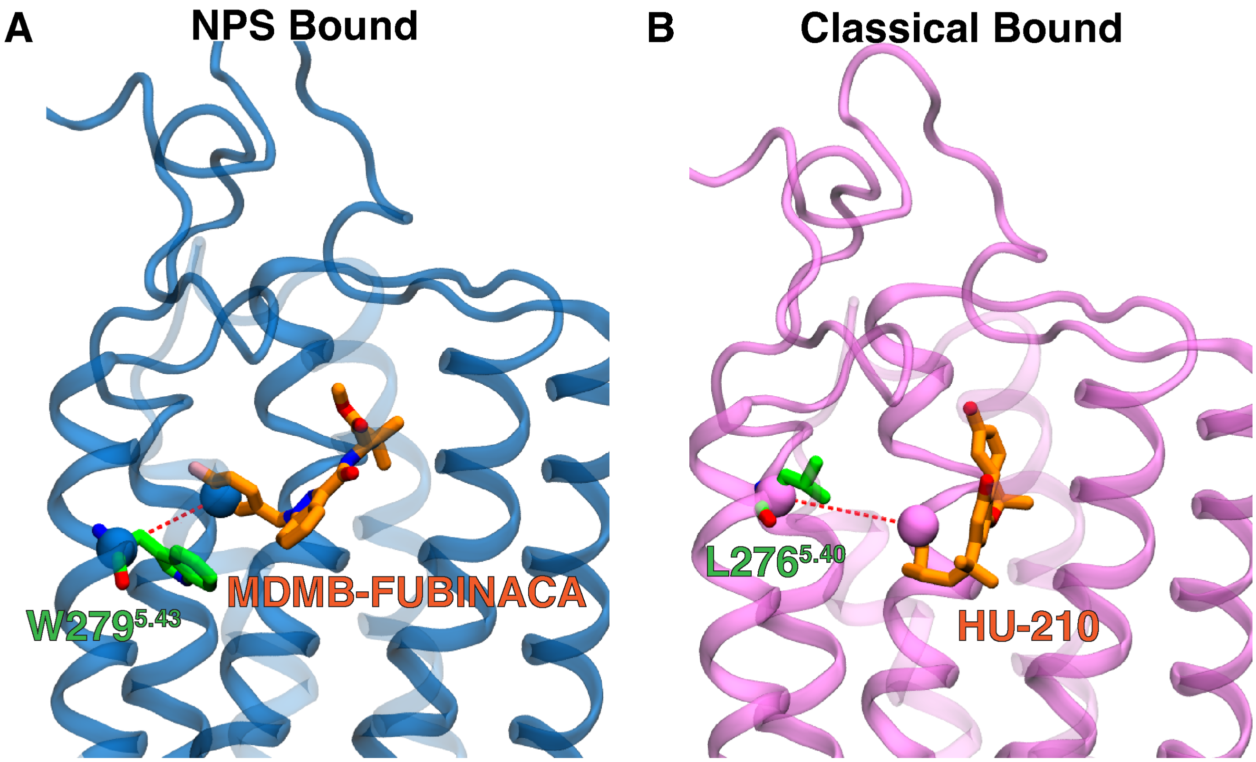


Figure S14: Modeled $CB_1$ holo structures are shown as cartoon representation with the liigands (MDMB-FUBINACA (A) and HU-210 (B)) in the orthosteric bound pose. TM5 distances from the ligands are presented as red dotted line. Ligands and residues in TM5 are represented as sticks. Atoms of interest are shown as vdw representation.

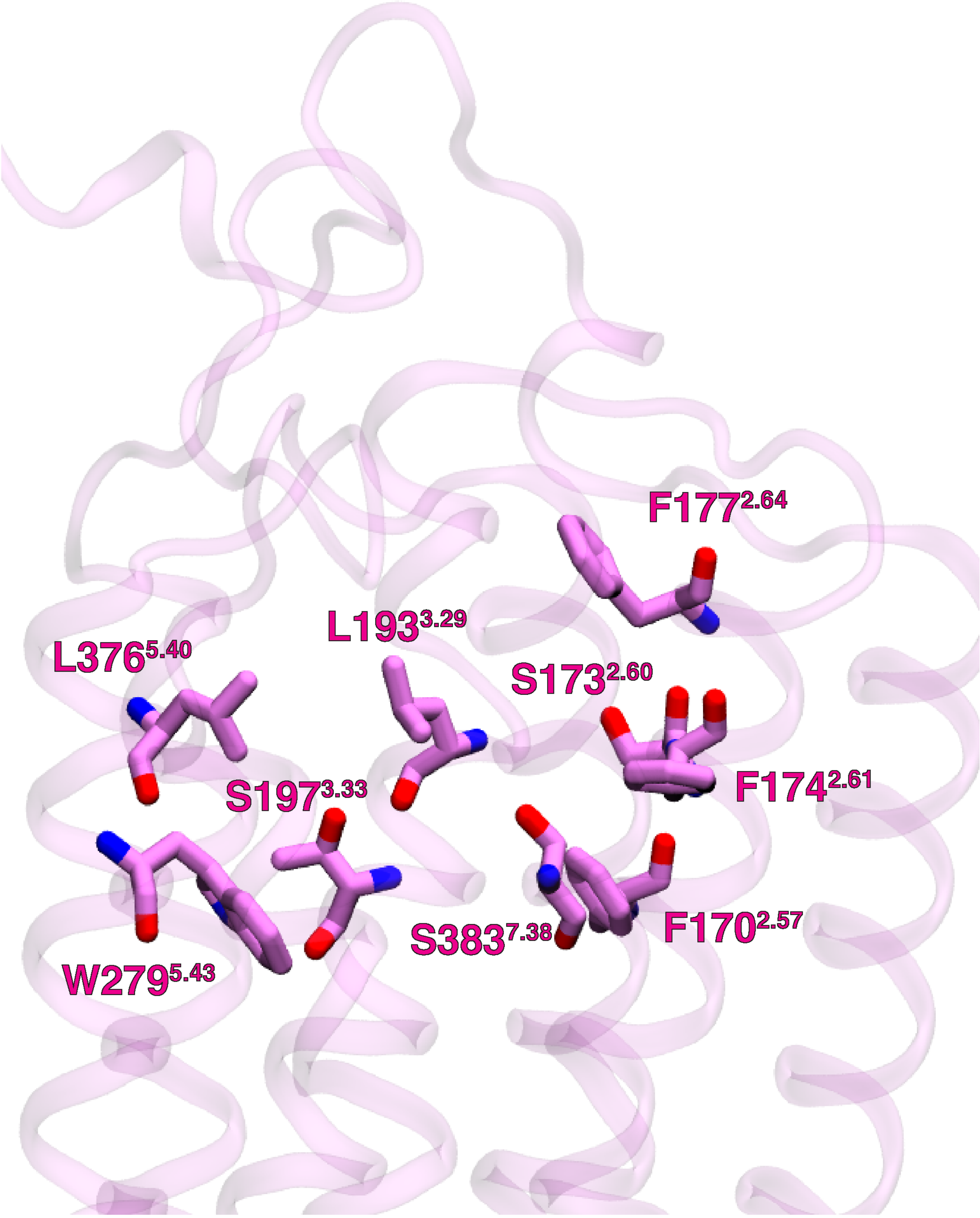


Figure S15: Binding pocket residues are highlighted as sticks. These residues are selected from TM2, TM3, TM5, TM7. These residues were considered for feature calculation to build MSM and TRAM.

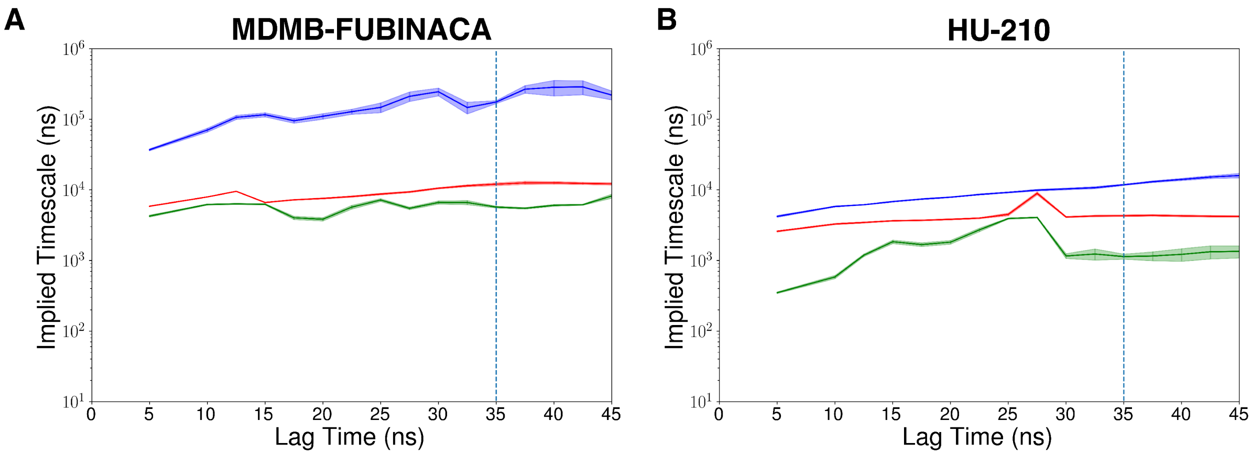


Figure S16: Top three implied timescales were plotted against different lag times for unbinding simulations of MDMB-FUBINACA (A) and HU-210 (B). MSM lagtimes were selected to be 35 ns for both ligands. For MDMB-FUBINACA, these calculations were performed with 700 clusters and 7 tIC dimensions. For HU-210, these calculations were performed with 800 clusters and 6 tIC dimensions. Errors in the implied timescale were calculated using 3 bootstrapped samples, where each sample containing 95% of the original unbiased data.

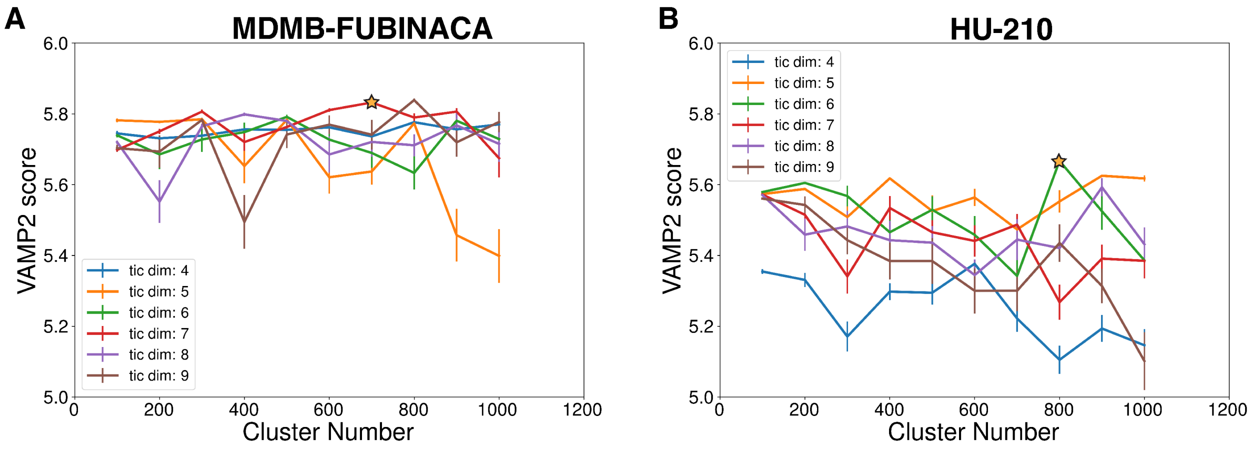


Figure S17: VAMP-2 scores of MSMs built with different cluster numbers are shown for MDMB-FUBINACA (A) and HU-210 (B) unbinding simulations. Different number of tICs used for MSM building were shown with different colors. The optimal VAMP-2 score in each case is marked with a star symbol.

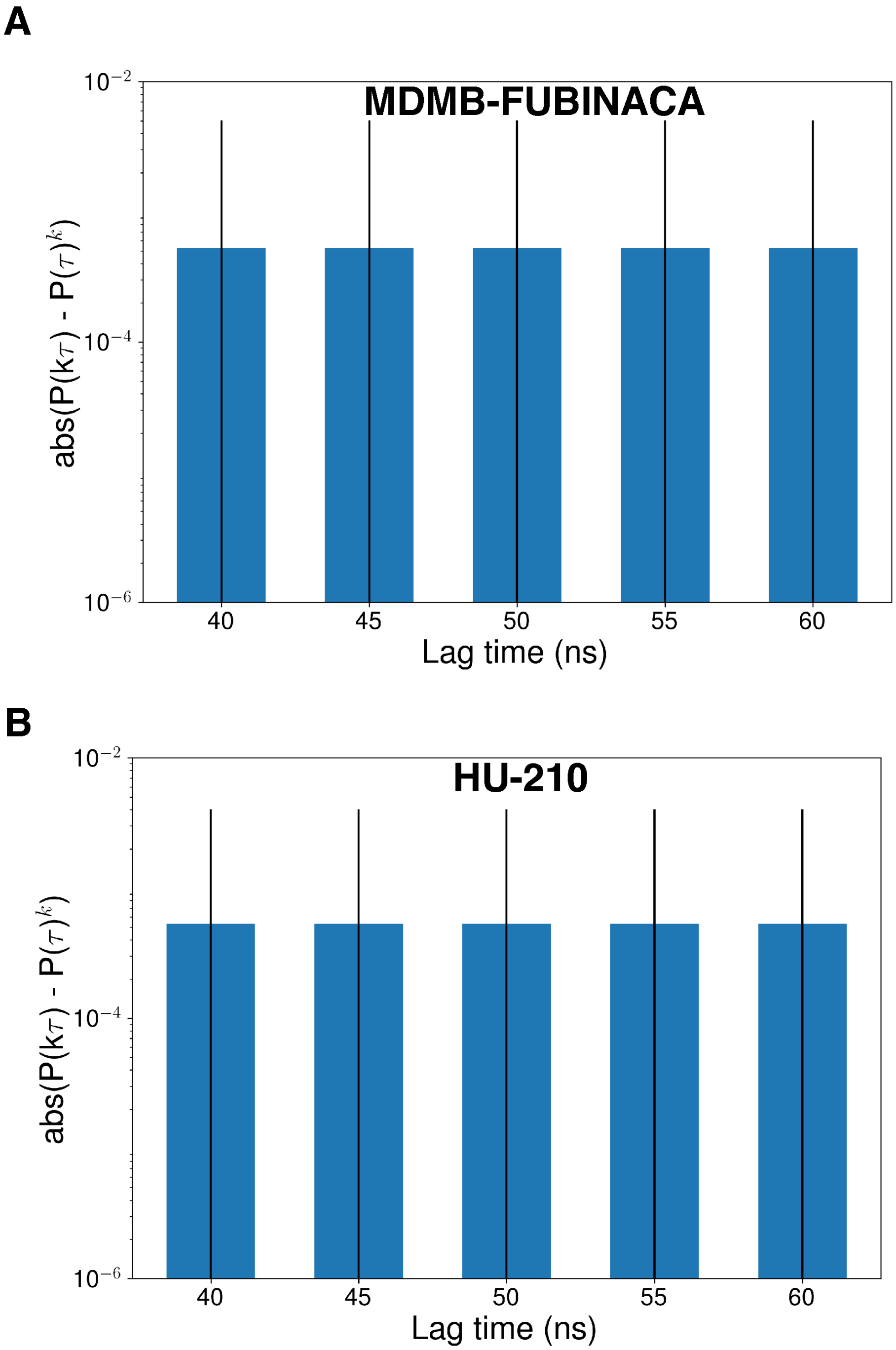


Figure S18: Absolute differences between $P(\tau)^k$ and $P(k\tau)$ are shown as bar plots for MDMB-FUBINACA (A) and HU-210 (B) unbinding simulations. Different values of k are considered where $\tau$ is 35 ns.

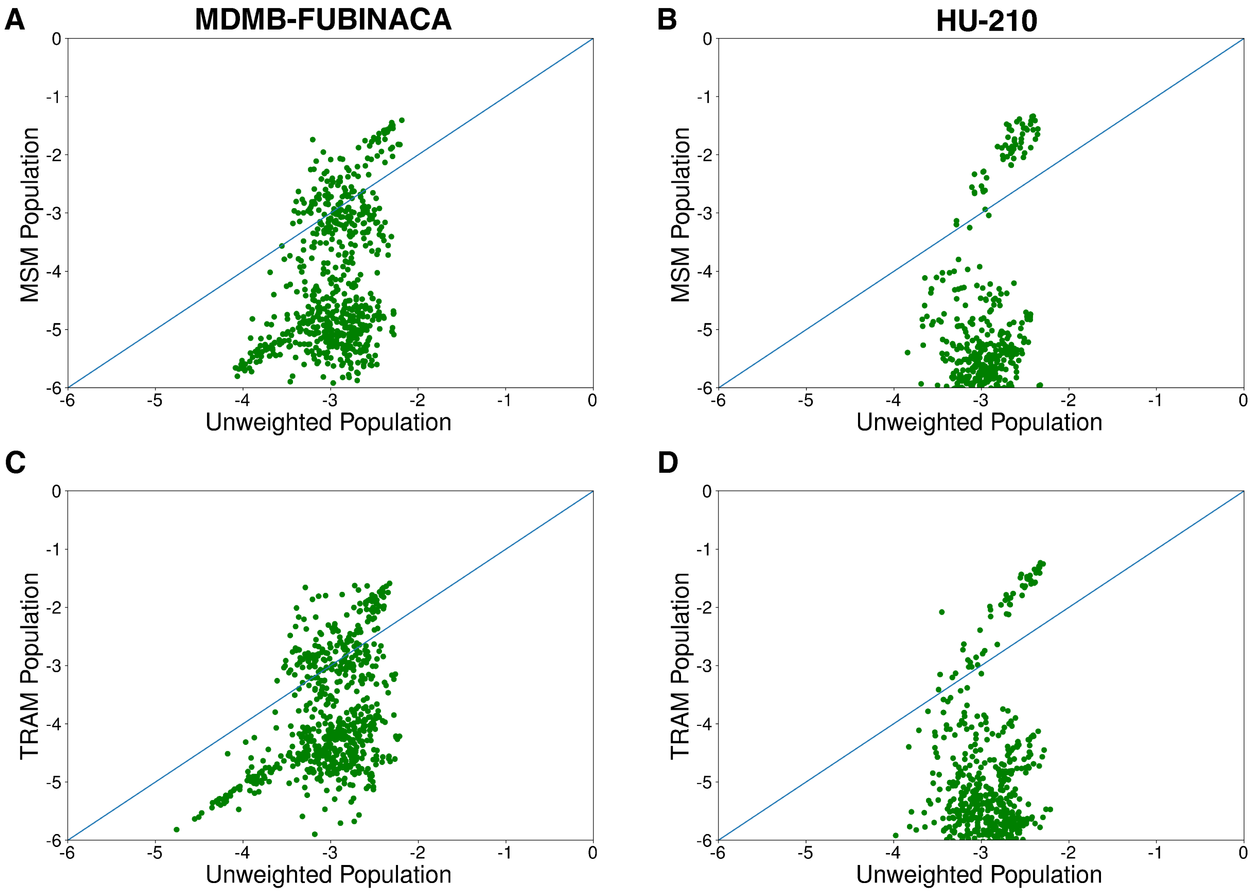


Figure S19: Raw probability and MSM weighted probabilities of the clustered states are plotted against each other for MDMB-FUBINACA (A) and HU-210 (B) unbinding simulations. Raw probability and TRAM weighted probabilities of the clustered states are plotted against each other for MDMB-FUBINACA (A) and HU-210 (B) unbinding simulations.

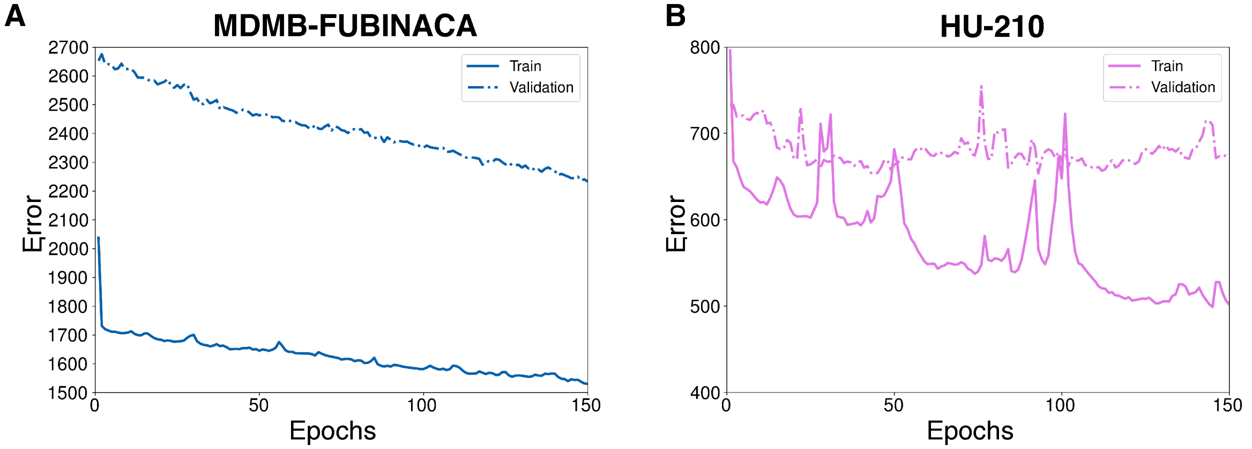


Figure S20: The reconstruction errors in the training (solid line) and validation (dotted line) data after per epoch training of NRI network for MDMB-FUBINACA (A) and HU-210 (B) bound trajectories.
